# Bioluminescent reporter influenza A viruses to track viral infections

**DOI:** 10.1101/2025.07.15.664884

**Authors:** Ramya S. Barre, Ahmed Mostafa, Kevin Chiem, Rebecca L. Pearl, Roy N. Platt, Anastasija Cupic, Timothy J. C. Anderson, Ulla G. Knaus, Randy A. Albrecht, Adolfo García-Sastre, James J. Kobie, Aitor Nogales, Luis Martinez-Sobrido

**Affiliations:** Texas Biomedical Research Institute, San Antonio, TX, USA; Department of Microbiology, Immunology, and Molecular Genetics, University of Texas Health Sciences Center at San Antonio, San Antonio, TX, USA; Center of Scientific Excellence for Influenza Viruses, National Research Centre, Giza, Egypt; Department of Microbiology, Icahn School of Medicine at Mount Sinai, New York, NY, USA; Global Health Emerging Pathogens Institute, Icahn School of Medicine at Mount Sinai, New York, NY, USA; Graduate School of Biomedical Sciences, Icahn School of Medicine at Mount Sinai, New York, NY, USA; Conway Institute, School of Medicine, University College Dublin, Dublin 4, Ireland; Department of Medicine, Division of Infectious Diseases, Icahn School of Medicine at Mount Sinai, New York, NY, USA; Tisch Cancer Institute, Icahn School of Medicine at Mount Sinai, New York, NY, USA; Department of Pathology, Molecular and Cell-Based Medicine, Icahn School of Medicine at Mount Sinai, New York, NY, USA; Icahn Genomics Institute, Icahn School of Medicine at Mount Sinai, New York, USA; Heersink School of Medicine, Infectious Diseases, University of Alabama at Birmingham, Birmingham, Alabama, AL, USA; Center for Animal Health Research, CISA-INIA-CSIC, Madrid, Spain

**Keywords:** Seasonal Influenza, bioluminescent reporter, nanoluciferase, *in vivo* imaging systems, H1N1, non-structural segment

## Abstract

Influenza A viruses (IAV) infect a wide range of mammal and bird species and are responsible for seasonal outbreaks and occasional pandemics. Studying IAV requires methods to detect the presence of the virus in infected cells or animal models. Recombinant IAV expressing fluorescent proteins have allowed monitoring viral infection in cultured cells and *ex vivo* in the organs of infected animals. However, fluorescent-expressing IAV are often attenuated and are not suited for the imaging of infected animals using *in vivo* imaging systems (IVIS). To overcome this limitation, we generated a recombinant A/California/04/2009 H1N1 (pH1N1) expressing nanoluciferase (Nluc) from the non-structural (NS) viral segment (pH1N1-Nluc) that replicates efficiently *in vitro*, with growth kinetics and plaque morphology comparable to wild-type pH1N1 (pH1N1-WT). We used this pH1N1-Nluc to demonstrate its ability to identify neutralizing antibodies and antivirals, with neutralization and inhibition results comparable to pH1N1-WT. In mice, pH1N1-Nluc was able to induce similar body weight loss and mortality, and viral titers comparable to pH1N1-WT, results that were recapitulated in a ferret model of IAV infection. Using IVIS, pH1N1-Nluc enabled non-invasive, real-time tracking of viral infection *in vivo* and *ex vivo* following infection of mice with viral titers comparable to pH1N1-WT. The flexibility of this approach was further demonstrated by the generation of a Nluc-expressing recombinant A/Puerto Rico/8/1934 H1N1 (PR8-Nluc). Altogether, our results demonstrate that Nluc-expressing recombinant IAV represent a valuable tool for *in vitro* and *in vivo* studies, including the identification of antivirals and/or neutralizing antibodies, and to assess protective efficacy of vaccines.

**IMPORTANCE:** Despite the availability of recombinant influenza A viruses (IAV) expressing fluorescent reporter genes to track viral infections *in vitro* and *ex vivo*, these viruses are often attenuated and do not represent the best option for imaging entire animals using *in vivo* imaging systems (IVIS). To solve this limitation, we generated recombinant influenza pandemic A/California/04/2009 H1N1 expressing nanoluciferase (pH1N1-Nluc) from the viral non-structural (NS) segment and demonstrate how expression of Nluc does not affect viral replication *in vitro* or viral pathogenesis *in vivo.* Importantly, we demonstrate the feasibility of detecting pH1N1-Nluc infection *in vivo* using IVIS. We also validate the flexibility of this approach by generating an influenza A/Puerto Rico/8/1934 H1N1 (PR8-Nluc). Our results support the feasibility of using these recombinant IAVs expressing Nluc from the NS segment for *in vitro* and *in vivo* studies, including the identification of neutralizing antibodies and/or antivirals, and to assess protective efficacy of vaccines.

## INTRODUCTION

Influenza viruses are respiratory pathogens of the family *Orthomyxoviridae*. Influenza viruses are classified into four different types: influenza A virus (IAV), influenza B virus (IBV), influenza C virus (ICV), and influenza D virus (IDV) (1). IAV and IBV are currently circulating in humans and responsible of seasonal infections that are associated with significant public health concerns and economic losses (1). IAV has also been responsible for causing pandemics of significant consequences to humans (2).

IAV is an enveloped virus with eight single-stranded negative sense viral segments, namely polymerase basic 2 (PB2), polymerase basic 1 (PB1), polymerase acidic (PA), hemagglutinin (HA), nucleoprotein (NP), neuraminidase (NA), matrix protein (M), and non-structural protein (NS) segments (1), each encoding one or more proteins (1). The smallest segment, NS, encodes at least two viral proteins: the nonstructural 1 (NS1) protein and the nuclear export protein (NEP), via mRNA splicing (1). The NS1 is a non-structural viral protein that plays a major role in virus replication and pathogenesis as an interferon (IFN)-antagonistic protein. NEP is involved in the export of viral RNAs from the host cell nucleus to the cytoplasm during viral infection (12).

In April 2009, a novel H1N1 IAV appeared in Mexico and the United States (3). This virus quickly spread worldwide, leading the World Health Organization (WHO) to declare it as the first influenza pandemic of the 21^st^ century in June 2009 and the third influenza pandemic involving an IAV H1N1 subtype (1, 4). Since then, this pandemic H1N1 (pH1N1) IAV has become seasonal in the human population. This event highlights the importance of H1N1 IAV infections in human health and the recurring potential of IAV H1N1 to cause pandemics in humans.

To easily track IAV infections *in vitro* and *in vivo*/*ex vivo* in animal models, fluorescent or bioluminescent viruses were developed by modifying viral segments to introduce one or more reporter gene(s) (5-9). We and others have previously show how IAV expressing fluorescent and/or bioluminescent reporter genes from different viral segments, including PB2, PB1, PA, HA, NA, and NS represent a suitable option to track viral infection in cultured cells and validated animal models of infection (5-7, 10-19). In addition, we have also generated a bireporter IAV expressing a fluorescent reporter gene (Venus) from the NS segment and a bioluminescent nanoluciferase (Nluc) gene from the HA segment (8). One of the advantages of using recombinant viruses expressing fluorescent reporter genes is their ability to identify the presence of infected cells in cultured cells or *ex-vivo,* from tissues of infected animals using *in vivo* imaging systems (IVIS) (8, 20). However, recombinant IAV expressing fluorescent proteins have been shown to be attenuated both, *in vitro* and *in vivo* (9, 12, 21, 22). Although recombinant IAV expressing fluorescent reporter genes are suitable for *in vivo* imaging using IVIS, their utility is limited by the size or body mass of the infected animal and are frequently disturbed by tissue autofluorescence, resulting in substantial background (23, 24). Moreover, it has been shown that recombinant IAV expressing fluorescent proteins could easily lose reporter expression after a few passages of the virus in cultured cells (9, 21, 25, 26). Bioluminescence (e.g., Nluc) offers the advantage of not affecting viral replication in cultured cells or in animal models and the feasibility of tracking viral infections in entire animals using IVIS (27). Thus, recombinant IAV expressing Nluc have the advantage over fluorescent recombinant viruses to track viral infections, in the entire animals using IVIS (8, 10, 20, 28). Several recombinant IAV expressing Nluc from the viral polymerases have been previously described in the literature. However, to date, few recombinant IAV expressing Nluc from the viral NS segment has been described for avian H9N2 (18), PR8 (19) or PR8-based H5N1 and PR8-based H7N9 (29). Expression of Nluc from the NS segment offers several advantages. These include minimal impact on viral fitness, stable reporter expression over multiple passages, and compatibility with sensitive, non-destructive luminescence-based assays (19). Notably, none of these studies have been conducted in a pandemic H1N1 background.

In this study, we generated a recombinant influenza A/California/04/2009 H1N1 (pH1N1) expressing Nluc (pH1N1-Nluc) by introducing the Nluc open reading frame (ORF) fused to the C-terminal domain of the NS1 protein in a modified NS viral segment. *In vitro*, pH1N1-Nluc replicates and generate plaques of comparable size to pH1N1-WT. Importantly, we demonstrated that pH1N1-Nluc can be used to easily identify neutralizing monoclonal antibodies (MAbs) or antivirals with neutralization and inhibition titers, respectively, similar to those obtained with pH1N1-WT, opening the feasibility of using pH1N1-Nluc to interrogate large libraries of compounds in high-throughput screening (HTS) settings to identify neutralizing MAbs and/or antivirals. *In vivo,* pH1N1-Nluc has similar morbidity and mortality than pH1N1-WT but offers the opportunity to identify the presence of the virus in infected mice using IVIS. Notably, viral titers of pH1N1-Nluc in the nasal turbinate (NT) and lungs of infected animals were comparable to those in pH1N1-WT-infected mice, demonstrating the feasibility of using pH1N1-Nluc to evaluate viral pathogenicity, replication, and tissue tropism. We also evaluated viral infection and transmission of pH1N1-Nluc in ferrets and the feasibility of detecting the presence of the virus in tissues by Nluc signal. Importantly, pH1N1-Nluc shows genetic and phenotypic stability after serial passages in MDCK cells. The flexibility of this approach was further demonstrated by the generation of a Nluc-expressing recombinant influenza A/Puerto Rico/8/1934 H1N1 (PR8-Nluc). Altogether, our results demonstrate that Nluc-expressing recombinant IAV represent a valuable tool for *in vitro* and *in vivo* studies, including the identification of neutralizing MAbs and/or antivirals, and to assess protective efficacy of vaccines.

## RESULTS

### Generation and characterization of recombinant pH1N1-Nluc

To generate a replication-competent recombinant pH1N1 expressing Nluc, the Nluc ORF without the stop codon was cloned into the NS segment (**Fig 1**) as previously described (6, 28). The IAV NS segment encodes NS1 and NEP using an alternative splicing mechanism (**Fig 1A**). We first constructed a modified NS split (NSs) segment, where NS1 and NEP are expressed from a single transcript by using the porcine teschovirus-1 (PTV-1) 2A autoproteolytic cleavage, resulting in the expression of a NS1-2A-NEP transcript that is independently translated into NS1 and NEP (**Fig 1B**). Then, the Nluc ORF was cloned at the C-terminal of NS1 to generate the NSs plasmid encoding Nluc (NSs-Nluc) to produce the recombinant pH1N1-Nluc virus (**Fig 1B**). We next used our previously described plasmid-based reverse genetics approaches where the pH1N1 NS plasmid was substituted with the NSs-Nluc plasmid together with the other 7 plasmids encoding the rest of the pH1N1 viral segments to generate the pH1N1-Nluc virus.

**Figure 1.**
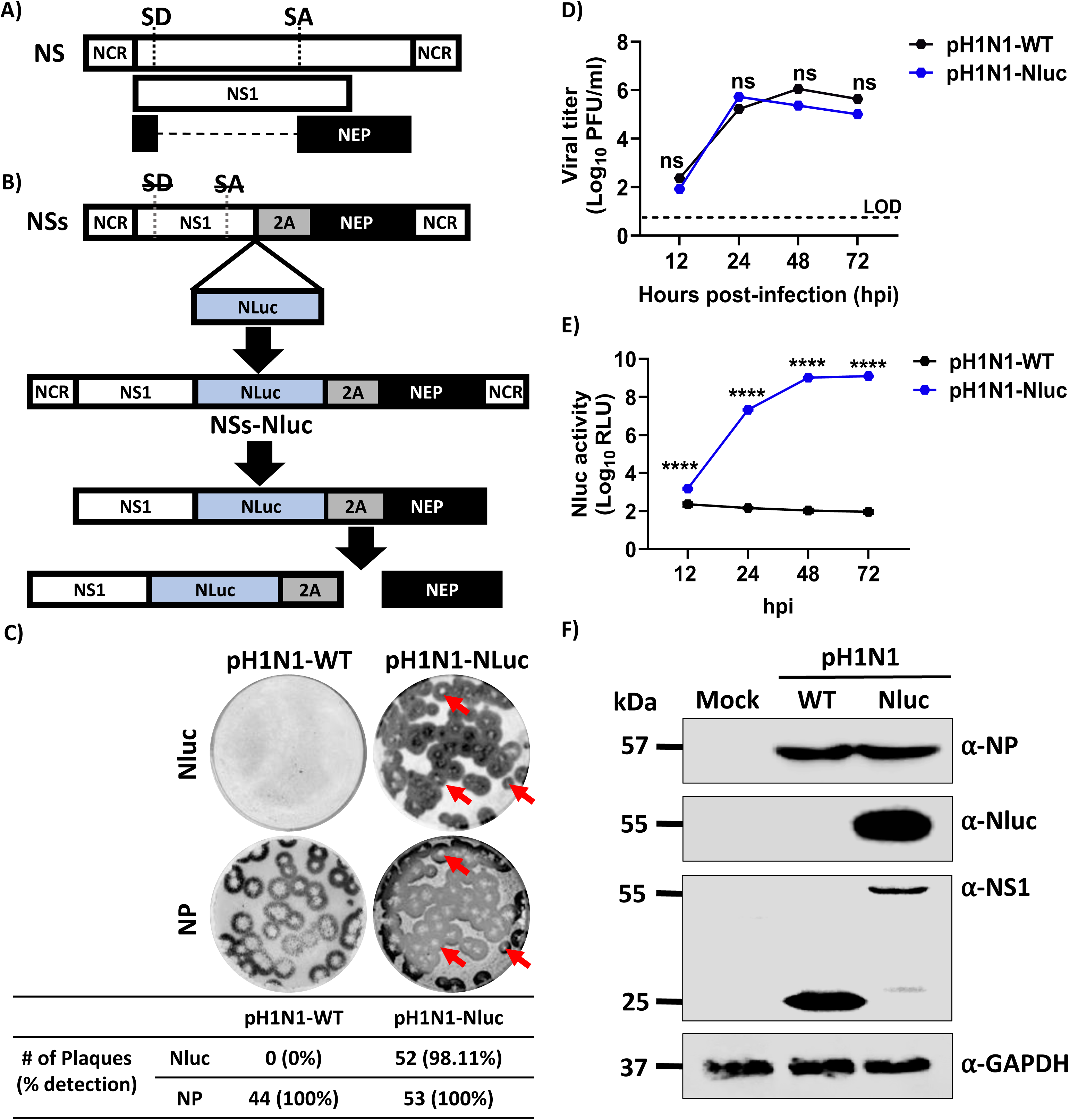
Generation and *in vitro* characterization of pH1N1-Nluc. A) **Schematic representation of NS viral segment.** The viral NS1 is represented in white and NEP in black. NCR, non-coding regions. **B**) **Schematic representation of NSs-Nluc viral segment encoding Nluc.** The Nluc coding sequence is represented in blue and the PTV-1 2A in a gray box. To inhibit the splicing process in the NSs constructs, silent mutations were introduced in the splice donor (SD, strikethrough) and acceptor (SA, strikethrough) sites. **C**) **Plaque phenotype of pH1N1-WT and pH1N1-Nluc in MDCK cells.** Viral plaques were evaluated at 72 hpi. Nluc staining (top) and NP immunostaining (bottom). Red arrows show the co-localization of Nluc staining (top) and viral plaques (bottom). **D**) **Multicycle growth kinetics of pH1N1-WT and pH1N1-Nluc viruses in MDCK cells.** Viral titers from culture supernatants of pH1N1-WT and pH1N1-Nluc infected (MOI 0.01) MDCK cells were determined using immunofocus assay at indicated times post-infection. Data represents means and SD for triplicates. The dotted line indicates the limit of detection (LOD) of the assay. **E**) **Nluc expression in culture supernatants from MDCK cells infected with pH1N1-WT- and pH1N1-Nluc.** Cell culture supernatants from the viral growth kinetics (D) were used to measure Nluc activity. **F**) **Western blots.** MDCK cells were infected (MOI=3) with pH1N1-WT or pH1N1-Nluc, or mock-infected. At 12 hpi, cell extracts were prepared, and Western blot was performed to assess levels of NP, NS1, and Nluc expression. Cellular GAPDH was used as a loading control. Data are represented as mean ± SD. A two-way repeated measure ANOVA with Geisser-Greenhouse correction. Post-hoc multiple comparisons performed using Šídák to compare groups within each time-point. The significant differences are indicated (ns=non-significant, **** = *p* < 0.0001).

After viral rescue, we next compared the plaque and replication phenotypes of pH1N1-Nluc to the wild-type pH1N1 (pH1N1-WT) (**Figs 1C and 1D**). Interestingly, the plaque phenotype of pH1N1-Nluc was comparable to pH1N1-WT (**Fig 1C**). Importantly, when viral plaques were stained with the anti-NP MAb HB-65, 98.11% of the viral plaques were Nluc positive by immunostaining (**Fig 1C**). When we evaluated viral growth kinetics, pH1N1-Nluc replicated to levels comparable to pH1N1-WT in MDCK cells at defined time points post-infection (**Fig 1D**). As expected, we were able to detect Nluc expression in the cell culture supernatants of MDCK cells infected with pH1N1-Nluc but not in cell culture supernatants of MDCK cells infected with pH1N1-WT (**Fig 1E**). We further demonstrated Nluc expression from cell extracts obtained from MDCK cells infected with pH1N1-Nluc by Western blot (**Fig 1F**). As expected and due to the carboxy-terminal extension with Nluc of the NS1 protein, we observed a higher molecular size of NS1 in MDCK cell extracts infected with pH1N1-Nluc compared to cells infected with pH1N1-WT (**Fig 1F**). Notably, levels of expression of viral NP were comparable in MDCK cells infected with pH1N1-Nluc or pH1N1-WT (**Fig 1F**). These results demonstrate that the recombinant pH1N1-Nluc has plaque and replication phenotypes that are similar to pH1N1-WT with the advantage to easily monitor viral infection by assessing Nluc expression from the cell culture supernatants of infected MDCK cells.

### Using pH1N1-Nluc to identify neutralizing MAbs and antivirals

Neutralizing MAbs and antiviral therapeutics are important in providing protection against IAV (30-32). Most IAV neutralizing and/or antiviral assays rely on detecting the presence of infected cells using secondary biochemical methods, which typically extend the duration of the assay (15, 33). To overcome this limitation, we evaluated the feasibility of using Nluc expression to identify neutralizing MAbs and antivirals against pH1N1 (**Fig 2**). To that end, we used a previously described neutralizing MAb (KPF1) (34, 35) and Ribavirin, a nucleoside analog previously shown to have antiviral activity against RNA and DNA viruses, including IAV (36-38). Importantly, we compared the neutralization and antivirals results using this Nluc-based microneutralization assay to previously described assays (34) and calculated the neutralization titer 50 (NT_50_) and the inhibitory concentration 50 (IC_50_) of KPF1 MAb and ribavirin, respectively. Moreover, we compared results obtained with pH1N1-Nluc to those obtained with pH1N1-WT. The NT_50_ of KPF1 using Nluc activity (NT_50_=3.096 ng/ml) (**Fig 2A**) was comparable to the neutralization activity of KPF1 using a conventional microneutralization assay based on percentage of infection achieved by pH1N1-Nluc (NT_50_=4.458 ng/ml) (**Fig 2C**) or by pH1N1-WT (NT_50_=4.953 ng/ml) (**Fig 2E**). Likewise, the IC_50_ of ribavirin using the pH1N1-Nluc in the Nluc-based microneutralization assay (IC_50_=1.953 ng/ml) (**Fig 2B**) was comparable to that obtained with pH1N1-Nluc (**Fig 2D**) or pH1N1-WT (**Fig 2F**) using the classical microneutralization assay (IC_50_=2.576 ng/mL and IC_50_=2.584 ng/ml, respectively). These results demonstrate the feasibility of using pH1N1-Nluc in a Nluc-based microneutralization assay to identify neutralizing MAbs or compounds with antiviral activity against pH1N1-WT with similar NT_50_ and IC_50_ values.

**Figure 2.**
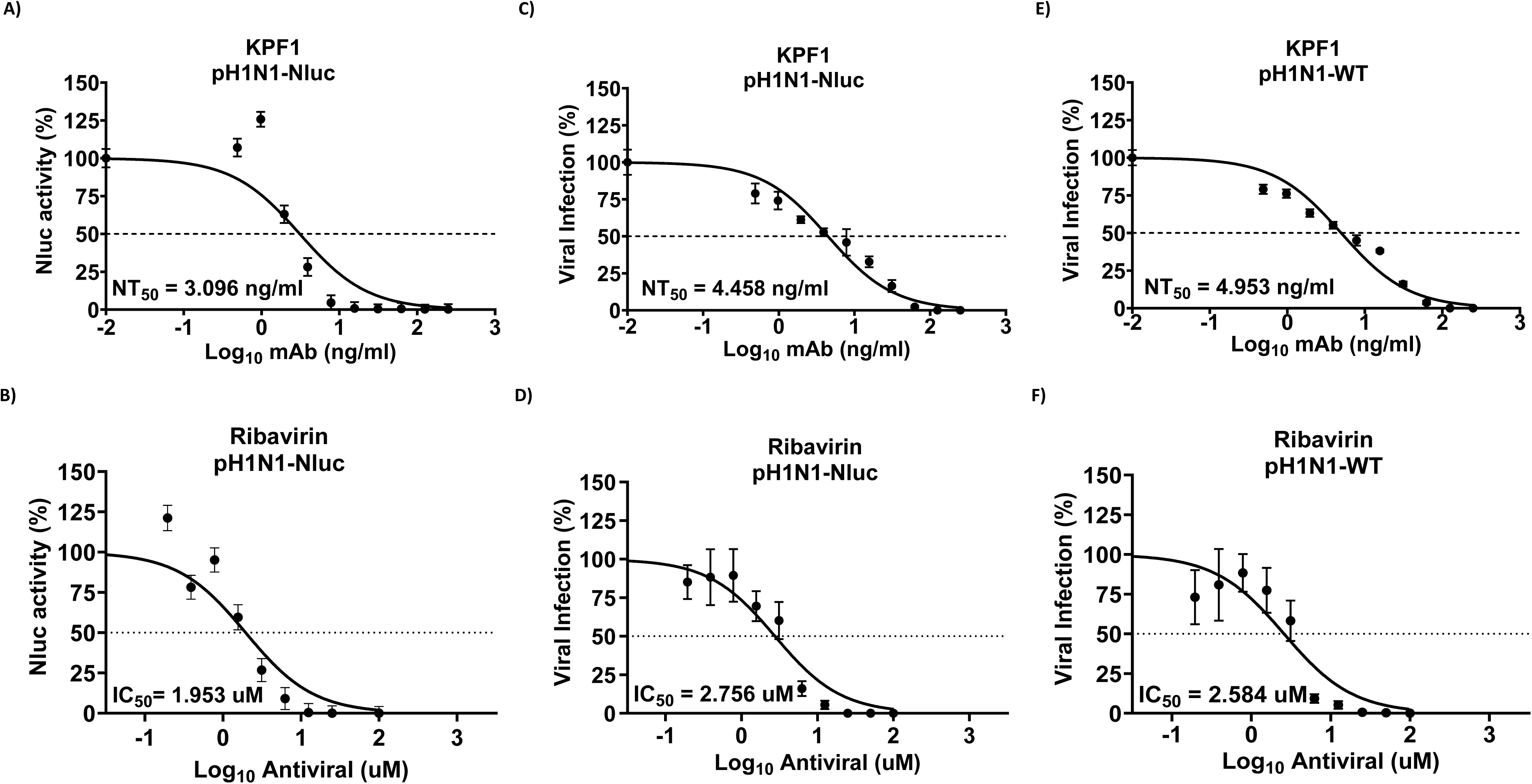
Microneutralization assays to evaluate neutralizing antibodies and antivirals. Monolayers of MDCK cells (96-well plate format, 10E4 cells/well, quadruplicates) were infected with 100 PFU of pH1N1-Nluc or pH1N1-WT virus for 1 h. After viral absorption, the virus inocula were removed, and cells were incubated with 2-fold serial dilutions of (**A**) a KPF1 pH1N1 neutralizing antibody (starting concentration of 0.25 µg/ml) or (**B**) Ribavirin (starting concentration of 100 µM). Virus neutralization (**A**) and inhibition (**B**) of pH1N1-Nluc was quantified by Nluc expression (left) using a luminometer or immunostaining titration of the viral titers (middle) at 45 hpi (left). Immunostaining was used to determine the viral titers in supernatants from pH1N1-WT-infected and treated samples (right). The 50% neutralization (NT_50_) of KPF1 and the 50% inhibitory concentration (IC_50_) of Ribavirin were calculated using sigmoidal dose-response curve. Dotted line indicates 50% neutralization (**A**) or inhibition (**B**).

### pH1N1-Nluc retains the virulence of pH1N1-WT in C57BL/6J mice

We next evaluated and compared the virulence of pH1N1-Nluc and pH1N1-WT in C57BL/6J (B6) mice (**Fig 3**). To evaluate morbidity and mortality, B6 mice (n=5) were infected, intranasally, with different doses (10^2^-10^5^ PFU) of pH1N1-Nluc (**Fig 3A**) or pH1N1-WT (**Fig 3B**) and mice were monitored for changes in body weight and survival. All mice infected with 10 PFU of pH1N1-Nluc or pH1N1-WT rapidly lose weight and succumbed to viral infection (**Figs 3A and 3B**). All B6 mice infected with 10 PFU of pH1N1-Nluc also succumbed to infection, while one mouse survived in the group infected with the same 10 PFU dose of pH1N1-WT, although this mouse presented signs of infection (**Figs 3A and 3B**). At 10³ PFU, two out of five mice survived infection with pH1N1-Nluc, while only one mouse survived in the pH1N1-WT group (**Figs 3A and 3B**). When infected with 10² PFU, two mice survived infection with pH1N1-Nluc, and three mice survived infection with pH1N1-WT (**Figs 3A and 3B**). The median lethal dose 50 (MLD_₅₀_) values were ∼2.89x10² PFU for pH1N1-Nluc and ∼1.67x10² PFU for pH1N1-WT (**Fig 3B**). These findings indicate that pH1N1-Nluc retains the virulence and lethality of pH1N1-WT in B6 mice.

**Figure 3.**
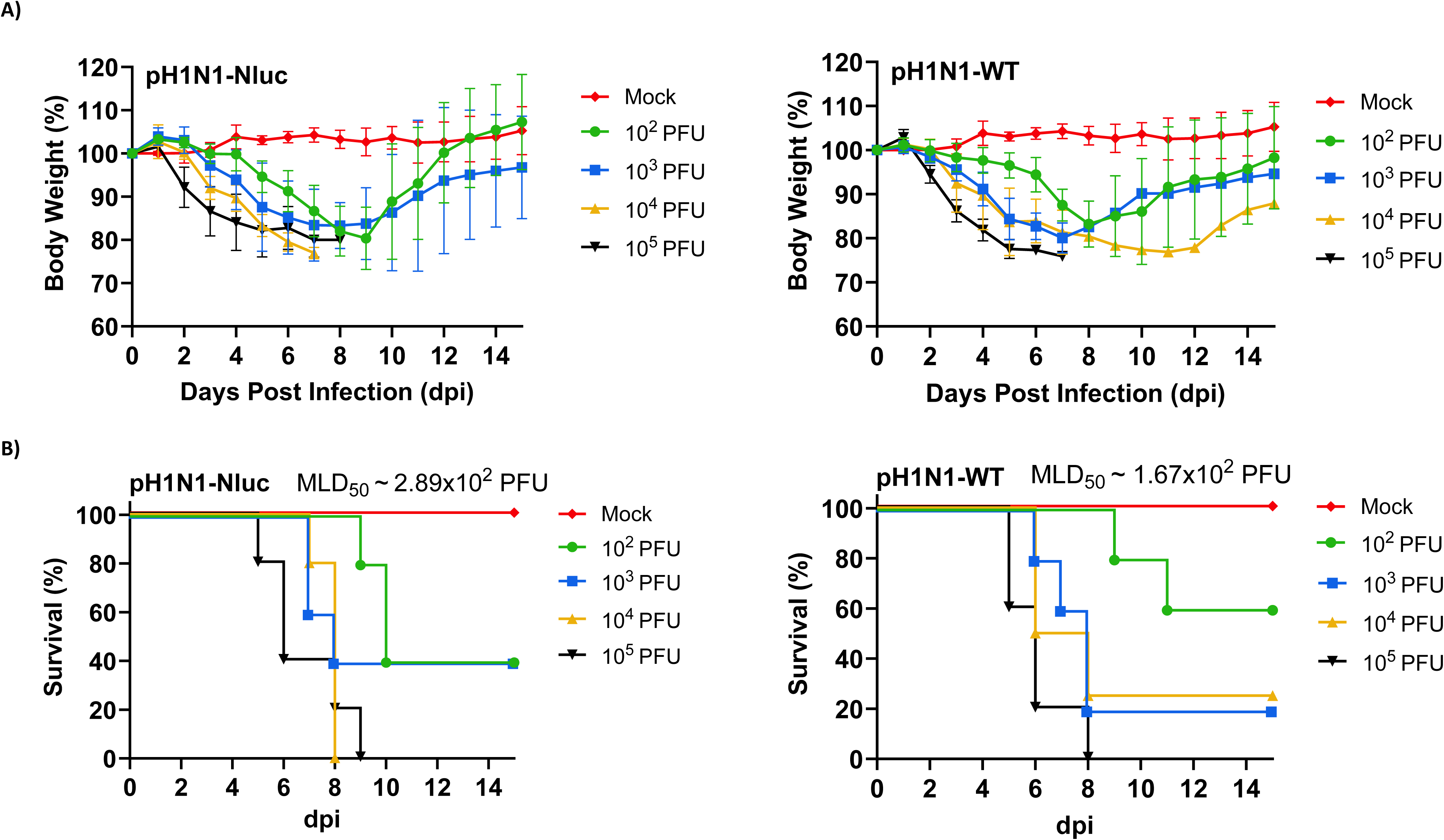
Pathogenicity of pH1N1-Nluc in mice. Female 6-week-old C57BL/6J mice (n=5) were inoculated with 10^2^, 10^3^, 10^4^, and 10^5^ PFU of pH1N1-Nluc or pH1N1-WT and monitored daily for 14 days. Mock-infected mice were used as controls. **A**) Percentage of body weight changes in mock-, pH1N1-Nluc-, and pH1N1-WT-infected C57BL/6J mice. **B**) Survival rates of mock-, pH1N1-Nluc-, and pH1N1-WT-infected C57BL/6J mice. Mice that lost 25% or greater of their initial weight were sacrificed. The MLD_50_ was calculated by Reed and Muench method. Data represents the means ± SD of the results for individual mice.

One of the advantages of using pH1N1-Nluc is the ability to track viral infection in living mice using *IVIS*. Consequently, mock-infected and pH1N1-Nluc- or pH1N1-WT-infected B6 mice were imaged with IVIS at 1, 2-, 4-, 6-, and 8-days post-infection (dpi) (**Fig 4**). We detected Nluc signal in mice infected with pH1N1-Nluc (**Fig 4A**). Nluc level was both dose and time dependent as we were able to detect an increase of the Nluc signal from day 1 to day 8; and mice infected with the highest dose showed higher levels of Nluc expression (**Fig 4A**). Notably, we were able to detect Nluc signals in the trachea of infected mice at early times post-infection (**Fig 4A**) that progress to the lungs at later times during viral infection (**Fig 4A**). Nluc level increased gradually but non-significantly from day 1 to day 8 in mice; with lowest levels of Nluc expression observed in mice infected with all doses of pH1N1-Nluc at day 1 post infection (**Fig 4C**). A dose- and time-dependent increase in Nluc luminescence was observed between 10^2^ PFU and 10^3-5^ PFU of pH1N1-Nluc, with no significant difference in luminescence beyond 10^3^ PFU, indicating a saturation effect at higher doses (**Fig 4A and 4C**). As expected, we were not able to detect Nluc expression in B6 mice infected with pH1N1-WT (**Figs 4B and 4D**).

**Figure 4.**
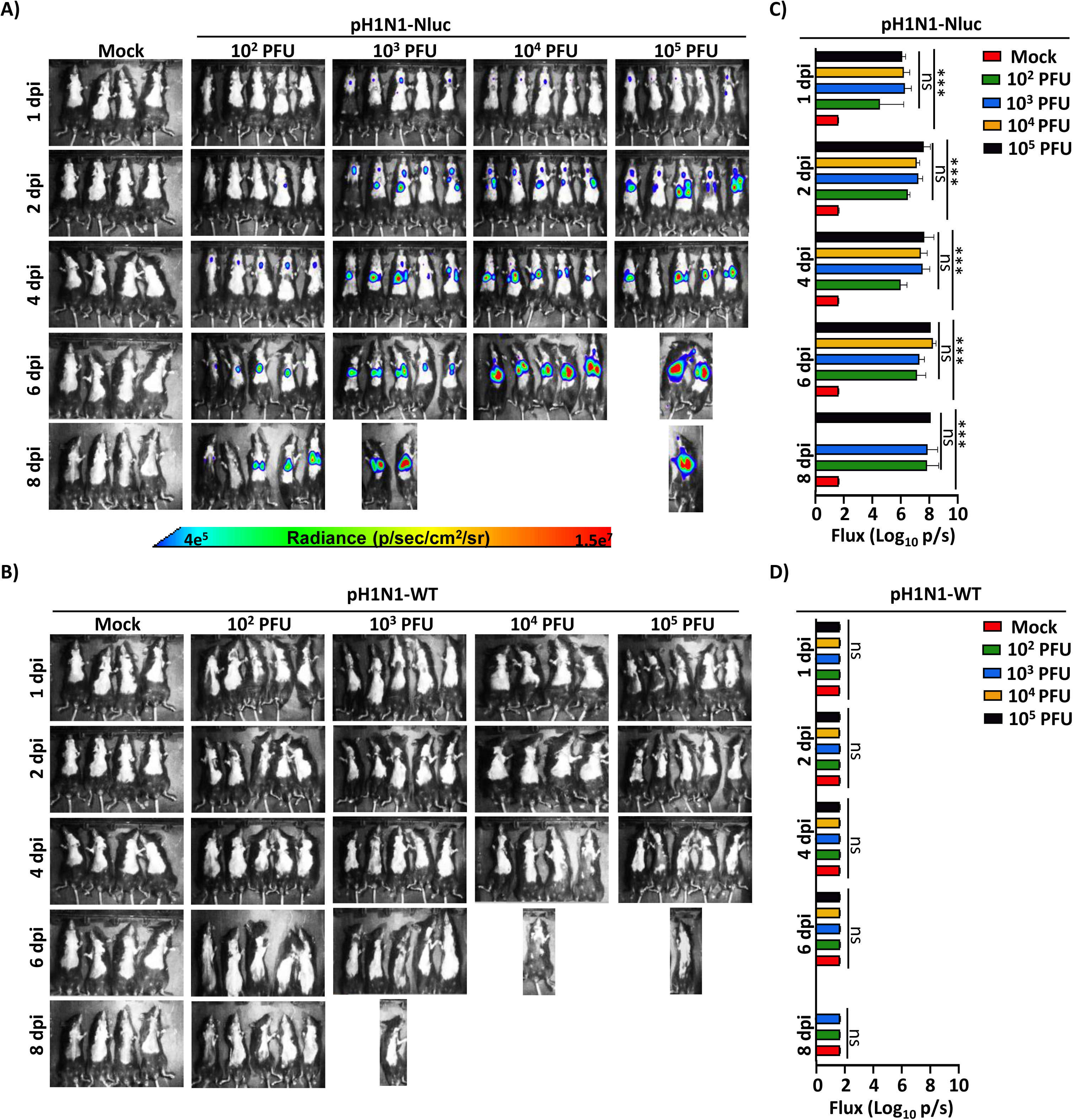
*In vivo* Nluc expression in pH1N1-Nluc-infected C57BL/6 mice. Female 6-week-old C57BL/6J mice infected as in Figure 3 were monitored for Nluc luminescence at 1, 2, 4, 6, 8 days post-infection (dpi) with pH1N1-Nluc (**A**) or pH1N1-WT (**B**) using IVIS. Radiance, defined as the number of photons per s per square cm per steradian (p s−1 cm−2 sr−1), is shown on the heat maps at the bottom. Quantification of Nluc luminescence in C57BL/6 mice infected with pH1N1-Nluc (**C**) or pH1N1-WT (**D**). A mixed-effects ANOVA followed by Dunnett’s multiple comparisons test was used (ns=non-significant, *** = *p* < 0.0001).

We next assessed Nluc level *ex vivo*, in the lungs of animals infected with pH1N1-Nluc (**Fig 5**). Briefly, different groups of mock-infected and pH1N1-Nluc- or pH1N1-WT-infected (10^2^-10^5^ PFU) B6 mice (n=6/group) were monitored for Nluc luminescence using IVIS before surgically excising the nasal turbinate (NT) and lungs on 2 and 4 dpi (n=3/group/day) (**Fig 5A**). Necropsied lungs were imaged *ex vivo* in the IVIS to detect luminescence (**Fig 5A**). Similar to results *in vivo* (**Fig 4**), Nluc luminescence in pH1N1-Nluc infected B6 mice was dose dependent (**Figs 5A and 5B**), with higher levels of Nluc expression observed in the lungs of mice infected with higher viral doses (10^3-5^ PFU) (**Figs 5A and 5C**). As expected, we were not able to detect Nluc expression in the excised lungs of mice infected with pH1N1-WT (**Figs 5B and 5D**).

**Figure 5.**
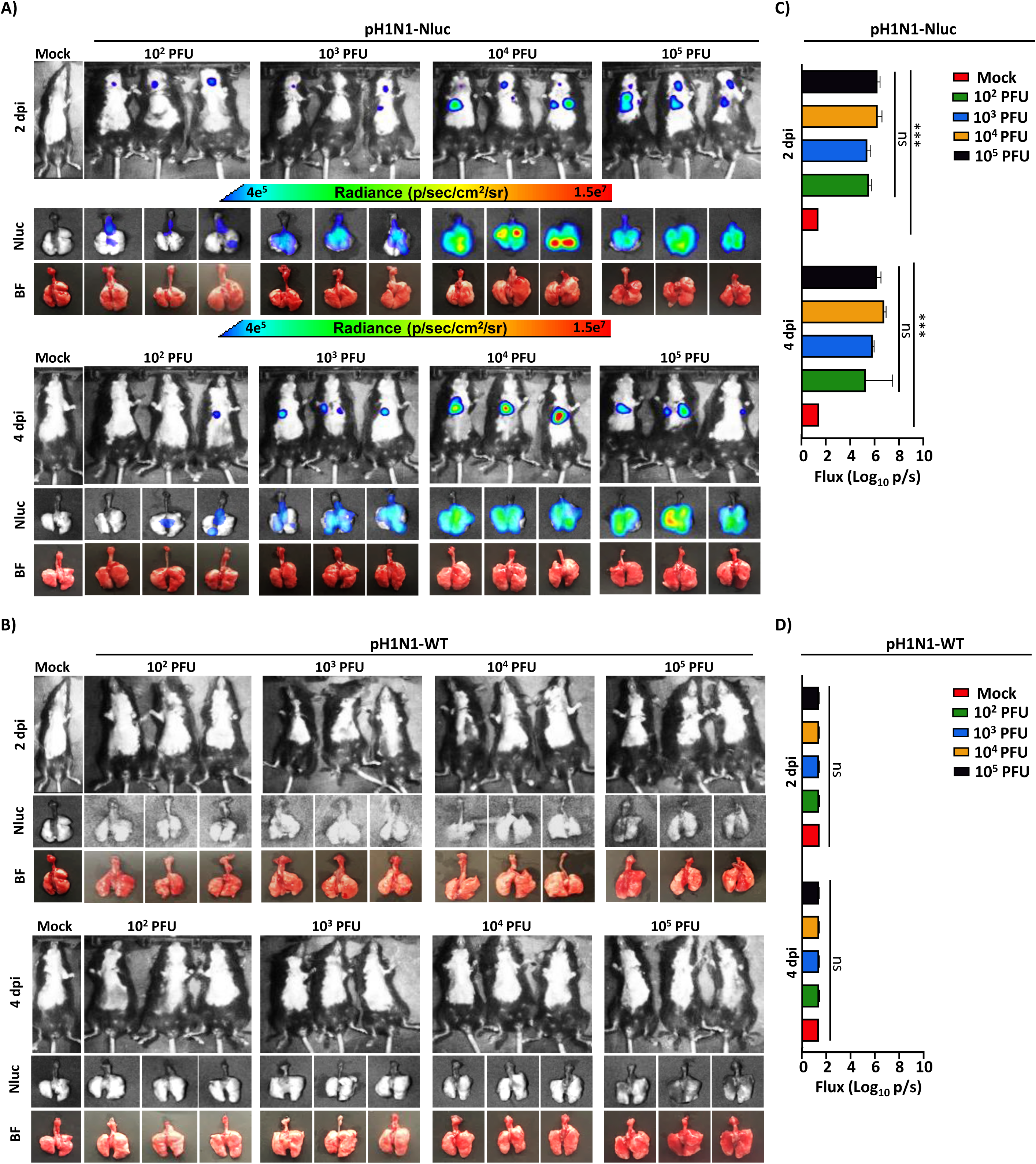
*In vivo* and *ex vivo* imaging of C57BL/6J mice infected with pH1N1-Nluc. **A**) Female 6-week-old C57BL/6J mice were infected with the indicated doses of pH1N1-Nluc. At 2 and 4 dpi, Nluc luminescence was detected using IVIS. **B**) Female 6-week-old C57BL/6J mice were infected with the indicated doses of pH1N1-WT and luminescence was monitored using IVIS at 2 and 4 dpi. Quantifications of Nluc luminescence in C57BL/6J mice infected with pH1N1-Nluc (**C**) or pH1N1-WT (**D**). Radiance, defined as the number of photons per s per square cm per steradian (p s−1 cm−2 sr−1), is shown on each of the indicated heat maps. A mixed-effects ANOVA followed by Dunnett’s multiple comparisons test was used (ns=non-significant, *** = *p* < 0.001).

A dose-dependent increase in viral titers was observed in the nasal turbinates (NT) of mice infected with pH1N1-Nluc (**Fig 6A**). Importantly, viral titers in the NT and lungs of B6 mice infected with pH1N1-Nluc were comparable to those detected in pH1N1-WT-infected B6 mice (**Fig 6A**), with no significant differences. Notably, we were able to detect Nluc signals in the NT and lungs of B6 mice infected with pH1N1-Nluc (**Fig 6B**). Nluc signals were dose dependent as we observed higher levels of luminescence in mice infected with the highest dose of pH1N1-Nluc (**Fig 6B**). We were not able to detect Nluc signals in the NT and lungs of B6 mice infected with pH1N1-WT (**Fig 6B**). Altogether, these results indicate that pH1N1-Nluc replicated in the NT and lungs of infected B6 mice at levels comparable to pH1N1-WT and Nluc luminescence can be used as a valid surrogate to evaluate the presence of the virus in these tissues (**Figs 6A and 6B**).

**Figure 6.**
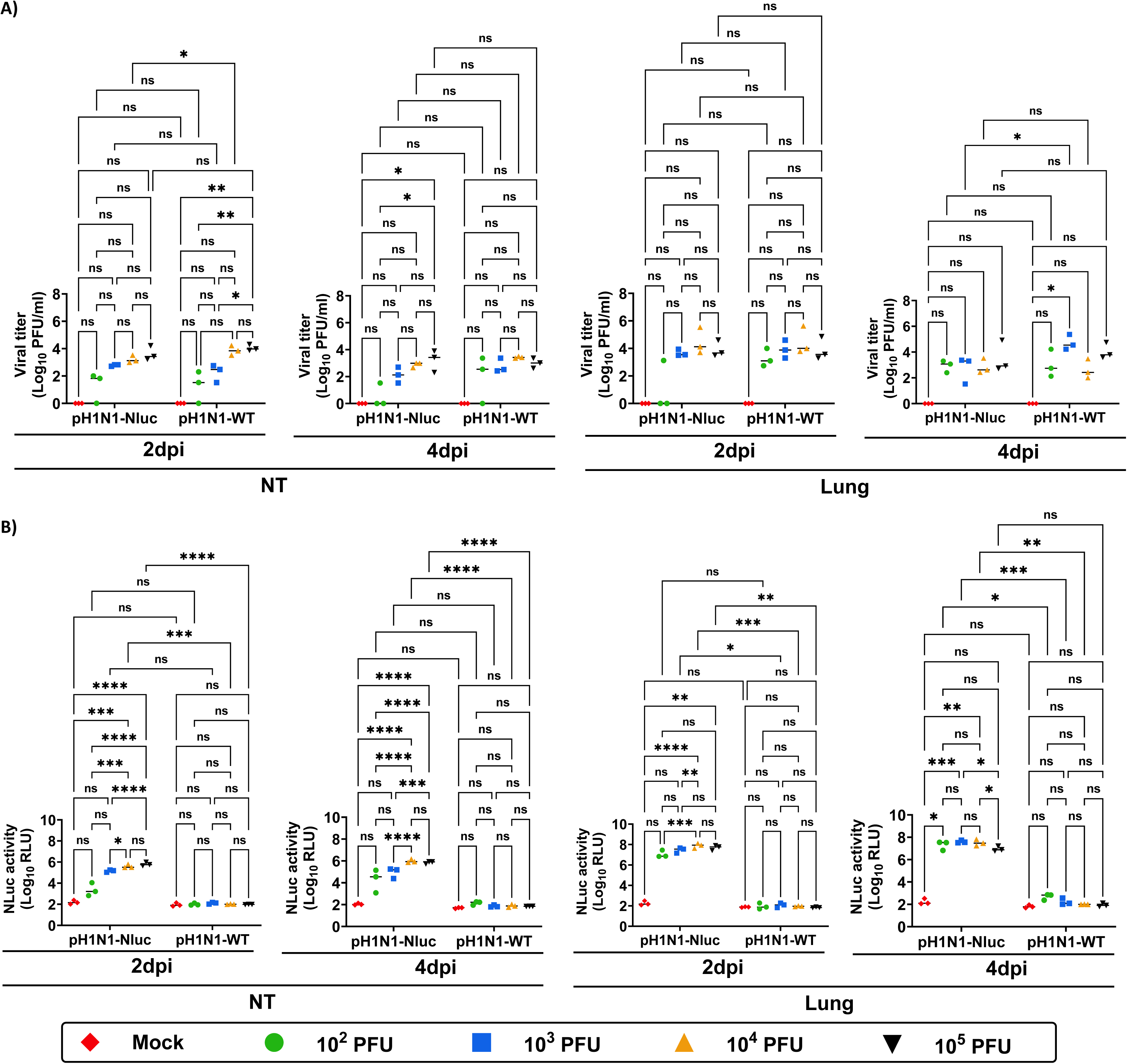
Viral titers and Nluc expression in C57BL/6J mice infected with pH1N1-Nluc. **A**) Viral titers in the nasal turbinate (NT) and lung tissues collected at 2 and 4 dpi are represented as Log_10_ PFU/ml. **B**) Nluc expression in tissue homogenates collected from the same NT and lung tissues of mice infected with pH1N1-Nluc and pH1N1-WT. Data are represented as mean ± SD. A mixed-effects ANOVA followed by Dunnett’s multiple comparisons test was used (ns=non-significant, * = *p* < 0.05, ** = *p* < 0.01, *** = *p* < 0.001, **** = *p* < 0.0001).

### Virulence and transmissibility of pH1N1-Nluc virus in ferrets

Ferrets have become one of the most used mammalian animal models to study influenza virus infection and transmission due to their susceptibility to IAV infection and similarity to humans in respiratory physiology and disease progression (39-41). To investigate the impact of the presence of Nluc in pH1N1-Nluc virus on viral virulence and transmissibility in ferrets, we infected separately 2 ferrets with 10^6^ PFU of pH1N1-Nluc or pH1N1-WT via intranasal route and caged individually before adding two naïve ferrets to each cage as contacts at 24 hpi (**Fig 7A**). Then, we measured body weight and collected nasal washes at 2, 4, and 6 dpi to determine viral titers in both infected and contact animals. Both pH1N1-WT (**Fig 7B**) and pH1N1-Nluc (**Fig 7C**) induced comparable levels of body weight loss (<15%) in experimentally infected ferrets albeit no body weight losses in contact animals up to 7 dpi. In ferrets infected with pH1N1-WT (**Fig 7D**) or pH1N1-Nluc (**Fig 7E**), viral titers in nasal washes increased from 2 to 4 dpi and were undetectable by 6 dpi. In contact ferrets, viral titers also increased from 2 to 4 dpi and began to decline by 6 dpi (**Figs 7D and 7E**). We detected Nluc luminescence in the nasal washes of pH1N1-Nluc-infected and contact ferrets, with Nluc levels decreasing from 2 to 6 dpi in infected ferrets, and Nluc levels increasing from 2 to 6 dpi in contact ferrets (**Fig 7E**), which correlated with viral titers determined by plaque assay from the same nasal washes (**Fig 7E**). The Nluc signal was not detected in the nasal washes of the pH1N1-WT-infected ferrets (**Fig 7D**).

**Figure 7.**
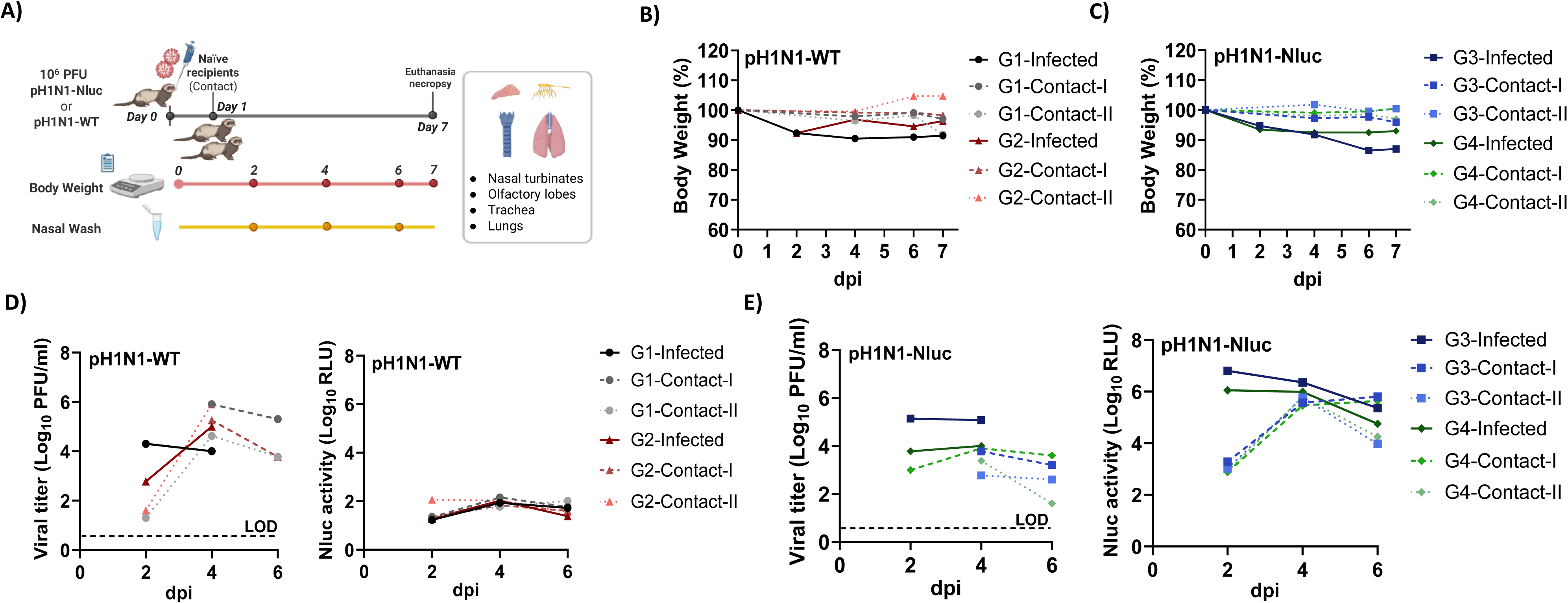
pH1N1-Nluc replication and transmission in ferrets. **A**) Experimental design of the ferret experiment. **B and C)** Percentages of body weight of ferrets infected (n=2) with pH1N1-WT (**B**) and pH1N1-Nluc (**C**) and contact ferrets (*n*=4). **D and E)** Nasal washes from pH1N1-WT- and pH1N1-Nluc-infected ferrets and their direct contacts, respectively, were titrated by plaque assay (left) or directly assessed for Nluc activity (**right**). The dotted line indicates the limit of detection (LOD) of the assay.

To investigate virus tropism, the NT, olfactory lobe (OL), trachea, and lungs of infected and contact ferrets were collected at 7 dpi and homogenized to evaluate viral titers. At 7 dpi, no viral titers were detected in tissues from ferrets experimentally infected with pH1N1-WT and pH1N1-Nluc after recovery (**Fig 8A**). However, we were able to detect high viral titers in the tissues collected from contact animals, with the highest levels in NT (**Fig 8B**), indicating recent and active infection with the virus following contact transmission from the infected animals. Levels of pH1N1-Nluc in the different tissues in contact ferrets were slightly lower than those in pH1N1-WT contact ferrets (**Fig 8B**). Importantly, no Nluc signal was observed in pH1N1-WT-infected or their contact ferrets (**Figs 8C and 8D**). In contrast, we detect high levels of Nluc luminescence in tissues from both pH1N1-Nluc infected and contact ferrets, with the highest levels of Nluc expression in NT (**Figs 8C and 8D**). Nluc was detected in the tissues of experimentally infected animals on day 7 at a time when infectious viruses were not detectable, indicating either higher sensitivity of the Nluc assay, or higher stability of Nluc in tissues as compared to infectious viruses (**Figs 8A and 8B**, right panels). These results demonstrate that while pH1N1-Nluc virus was slightly attenuated compared to pH1N1-WT in ferrets, we were able to detect Nluc luminescence that correlates with viral infection in the tissues of both experimentally infected and contact animals.

**Figure 8.**
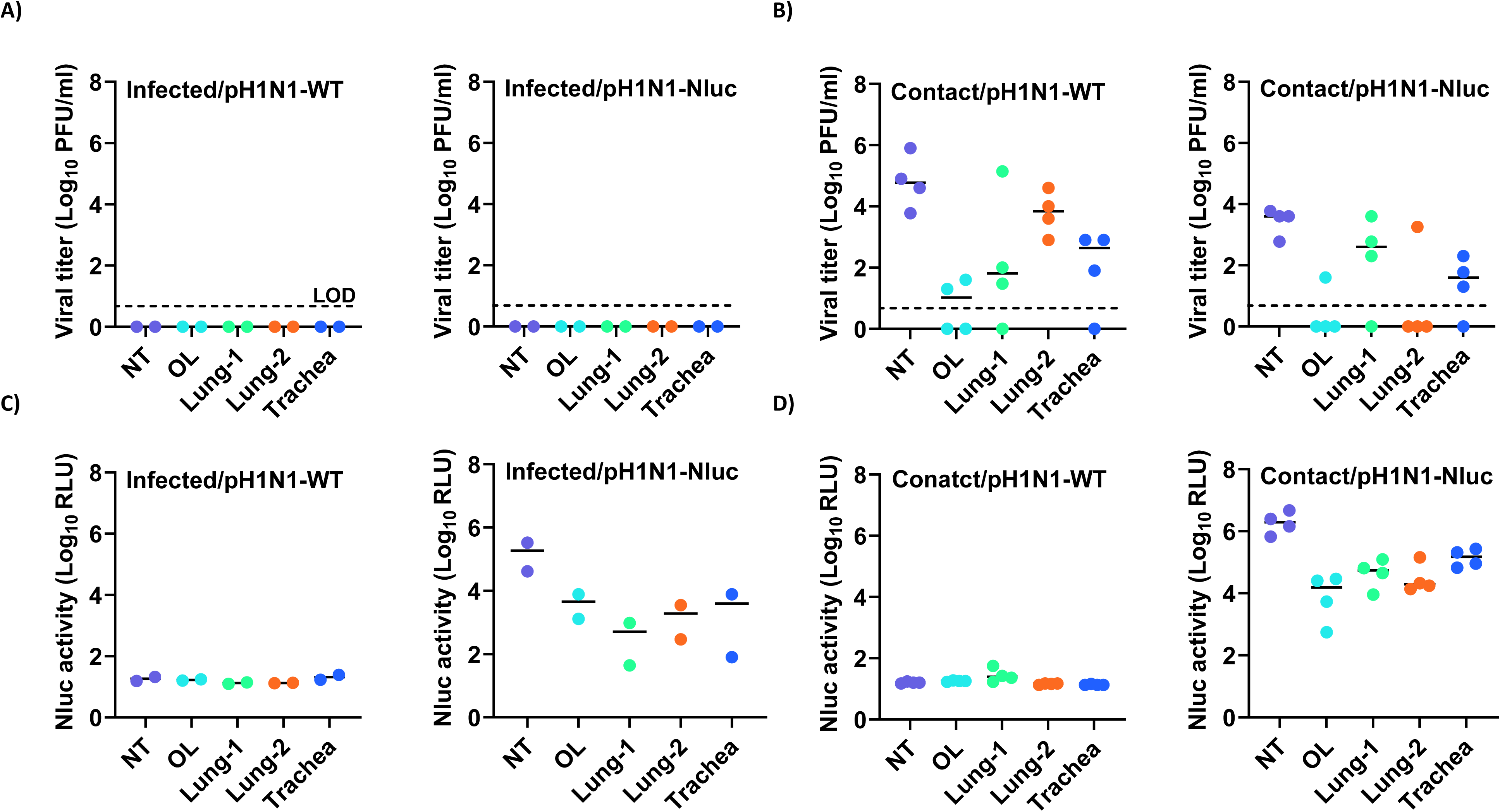
Viral titers and Nluc luminescence in pH1N1-Nluc- and pH1N1-WT-infected ferrets. (**A and B**) Viral titers in the nasal turbinate (NT), olfactory lobe (OL), lungs, and trachea infected (n=2) with pH1N1-WT and pH1N1-Nluc (**A**), and direct contact (*n*=4) animals (**B**), are represented as Log_10_ PFU/ml. (**C and D)** Nluc expressions in the NT, OL, lungs, and trachea of infected with pH1N1-Nluc and pH1N1-WT (**C**), and direct contact animals (**D**). The limit of detection (LOD) is indicated with a dashed line. Data are represented as mean ± SD.

### Genetic and phenotypic stability of pH1N1-Nluc

One important aspect for the use of reporter-expressing recombinant viruses is their genetic and phenotypic stability. Therefore, we next assessed the genetic and phenotypic stability of pH1N1-Nluc. To that end, pH1N1-Nluc was serially passaged in MDCK cells for 10 successive passages (**Fig 9**). Nluc expression was analyzed for passages P1-P10, showing stable Nluc expression over the 10 passages (**Fig 9A**). By plaque assay, pH1N1-Nluc virus collected at passage 1 (P1), passage 5 (P5), and passage 10 (P10) were positive for Nluc with Nluc expression reaching 97,56% (P1), 100% (P5), and 100% (P10) (**Fig 9B**). Importantly, NGS of pH1N1-Nluc collected from P1, P5, and P10 detected Nluc through the 10 serial passages (**Fig 9C**) with significant coverage (**Supplementary Table 1**). No significant Single Nucleotide Polymorphisms (SNPs) were detected in the NSs-Nluc viral segment (**Fig 9D**) and only a few point mutations were observed in other viral segments (**Supplementary Fig S1**). Contrary to previously described fluorescent-expressing recombinant IAV (9, 21, 25), this finding suggests that pH1N1-Nluc is genotypically and phenotypically stable.

**Figure 9.**
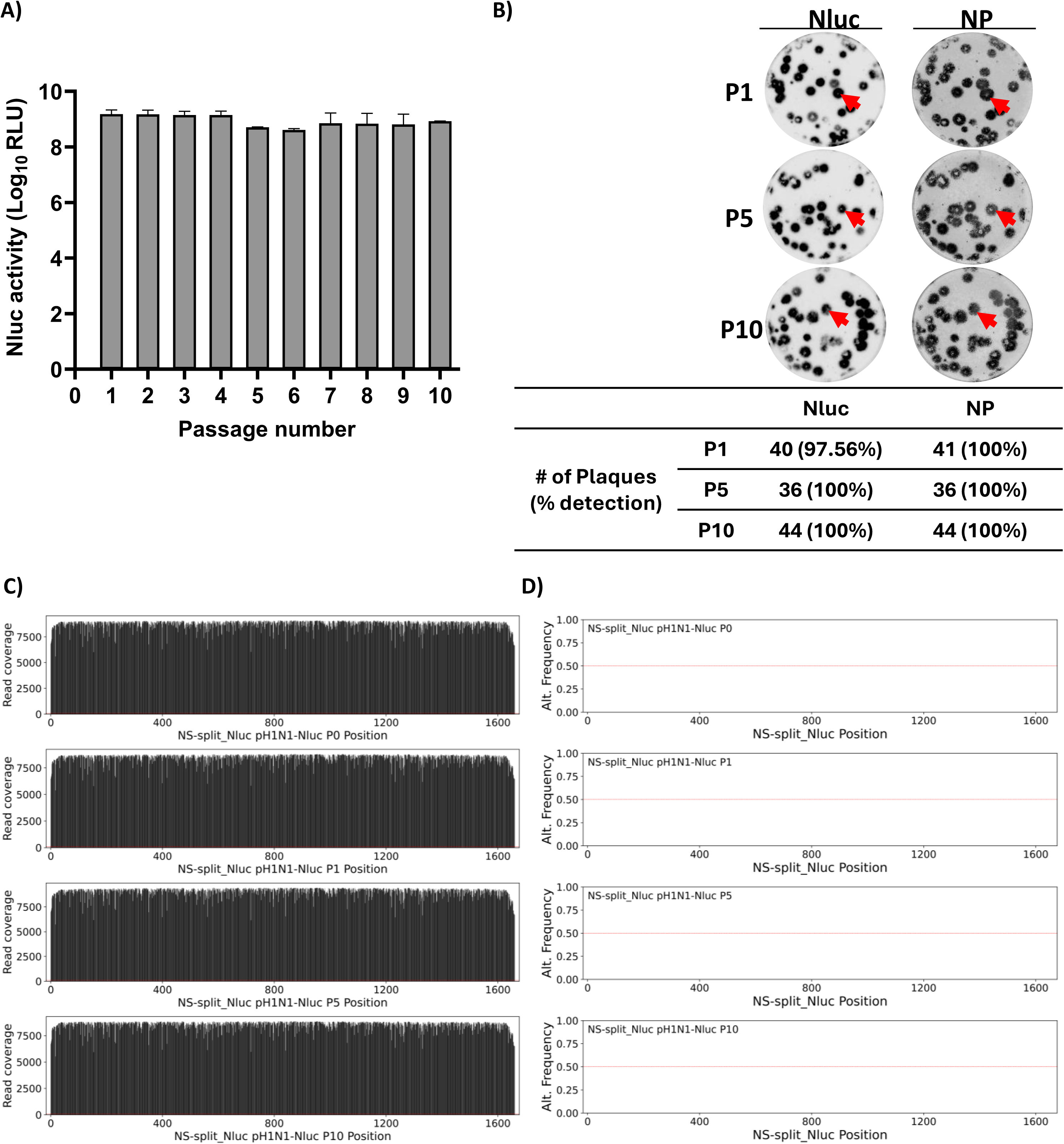
Stability of pH1N1-Nluc virus. MDCK cells were successively infected with pH1N1-Nluc for 10 serial passages. (**A**) Nluc expression in culture supernatants were measured for 10 viral passages. (**B**) Plaques of pH1N1-Nluc viruses, collected at passages 1 (P1), 5 (P5), and 10 (P10), were stained with Nluc substrate (left) or immunostained with a monoclonal antibody (HB-65) against the viral NP (right). **C**) Next generation sequencing (NGS) results covering the NSs-Nluc segment across passages 1, 5 and 10. **D**) Non-reference allele frequency - The passaged samples were compared to the sequence of used stock of recombinant pH1N1-Nluc virus (P0) to identify variants. The red line represents 50% allele frequency and would coincide with consensus sequence changes in the viral population.

### Generation and characterization of recombinant PR8-Nluc

To validate the feasibility of incorporating Nluc into the NS segment of a commonly used influenza A/Puerto Rico/8/1934 (H1N1) strain (PR8-WT), we genetically engineered and successfully rescued a recombinant PR8 virus expressing Nluc (PR8-Nluc) using similar experimental approaches to those described for pH1N1. The replication kinetics of PR8-Nluc were evaluated by plaque assay to assess viral growth dynamics and were compared to those of the parental PR8 (PR8-WT) (**Fig 10**). PR8-Nluc and PR8-WT showed comparable growth kinetics at 12-, 24-, 48- and 72 hpi (**Fig 10A**). The presence of Nluc in the same cell culture supernatants reflected a time-dependent increase of plateaued when viral titers remained stable (**Fig 10B**). As expected, Nluc activity in the cell culture supernatants of PR8-WT-infected MDCK cells was comparable to background levels (**Fig 10B**). In parallel, we also evaluated and compared the plaque morphology of PR8-Nluc to PR8-WT (**Fig 10C**). Plaque visualization using the Nluc substrate and immunostaining with the anti-NP HB-65 MAb revealed a comparable number (98.11%) and size of plaques for PR8-Nluc, demonstrating robust and consistent Nluc expression without impairing viral replication (**Fig 10C**). Additionally, Nluc expression was confirmed by Western blot using lysates of pH1N1-WT and pH1N1-Nluc infected MDCK cells (**Fig 10D**). These combined results support the feasibility of implementing the same approach for the generation of other H1N1 viruses for their use in *in vitro* and potentially *in vivo* studies to assess viral replication and to easily track viral infections.

**Figure 10.**
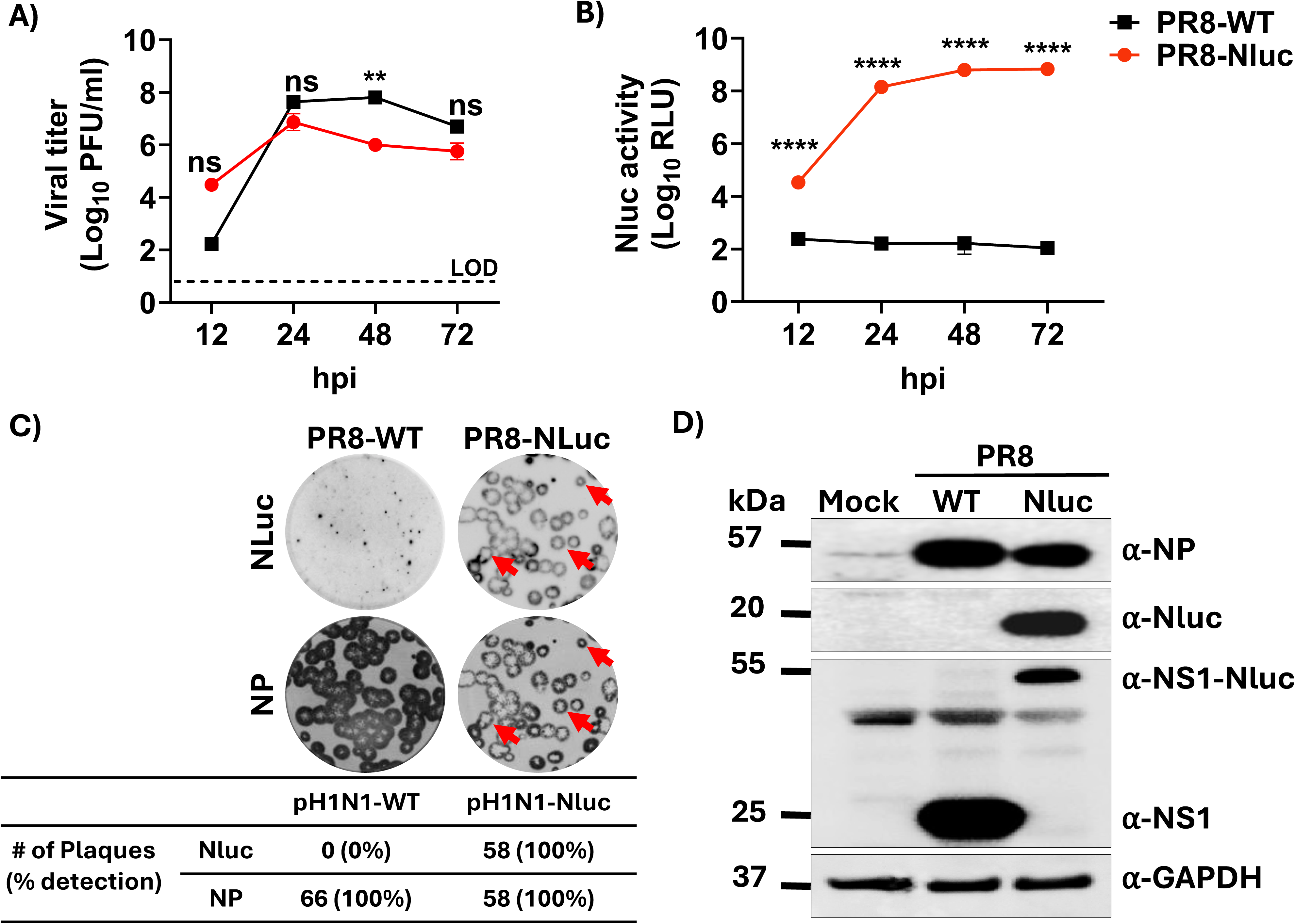
Generation and *in vitro* characterization of PR8-Nluc. (**A**) **Replication kinetics of PR8-WT and PR8-Nluc viruses in MDCK cells.** Viral titers from culture supernatants of PR8 and PR8-Nluc infected (MOI 0.01) MDCK cells were determined using immunofocus assay at 12, 24, 48 and 72 hpi. The dotted line indicates the limit of detection (LOD) of the assay. **B**) Nluc expression in culture supernatants at indicated time points in panel (A). Cell culture supernatants from the viral growth kinetics were used to measure Nluc activity. **C**) Plaque phenotype of PR8-WT and PR8-Nluc in MDCK cells. Viral plaques were evaluated at 72 hpi. Nluc staining (top) and NP immunostaining (bottom). Red arrows show the co-localization of Nluc staining (top) and viral plaques (bottom). **D**) Western blots. MDCK cells were infected (MOI=3) with PR8-WT or PR8-Nluc, or mock-infected. At 12 hpi, cell extracts were prepared, and Western blot was performed to assess levels of NP, NS1, and Nluc expression. Cellular GAPDH was used as a loading control. Data are represented as mean ± SD. A two-way repeated measure ANOVA with Geisser-Greenhouse correction was used. Post-hoc multiple comparisons were performed using Šídák to compare groups within each time-point. The significant differences are indicated (ns=non-significant, ** = *p* < 0.01, **** = *p* < 0.0001).

## DISCUSSION

Reporter-expressing IAV represent an excellent option for assessing viral replication, virulence, and transmission (7, 8, 11, 39, 42-44). In addition, recombinant IAV expressing fluorescent and/or luciferase reporter genes can be used to easily identify and/or characterize neutralizing MAbs, antiviral compounds, as well as to evaluate vaccine efficacy (12, 13). Fluorescent reporters tend to be unstable after multiple passages in cell culture and in several instances result in attenuation of the virus *in vitro* and *in vivo* (8, 22). Additionally, fluorescent proteins are not optimal for *in vivo* imaging of living animals using IVIS and can only be used to detect the presence of virus in infected tissues *ex vivo* (8, 20). In contrast, luciferase (e.g. Nluc) reporter genes offer several advantages over fluorescent proteins, including their small size, ATP-independent activity, high stability, and strong signal intensity, making them a more appropriated choice for imaging living animals using IVIS (11, 45, 46),

The NS segment of IAV, which encodes the NS1 protein and, via an alternative splicing mechanism, the NEP, has been frequently used for expressing reporter genes as fusions to the C-terminal domain of NS1 (8, 47). However, these studies have been limited to the expression of fluorescent proteins that often resulted in viral attenuation *in vitro* and *in vivo*. While previous studies have demonstrated the feasibility of generating recombinant IAV expressing Nluc from different viral segments (e.g., PB2, PB1, PA, HA, NS) (15-19), the feasibility of generating and detecting Nluc expression from recombinant seasonal H1N1 IAV from the NS segment had not been yet explored. Herein, we cloned Nluc fused to the C-terminal end of NS1 into a modified NS segment where NS1 and NEP are expressed from a single transcript. Using plasmid-based reverse genetics, we successfully generated a recombinant pH1N1-Nluc virus. *In vitro,* pH1N1-Nluc have comparable levels of viral replication and a similar plaque phenotype than pH1N1-WT. The advantages of pH1N1-Nluc expressing Nluc include ease of identifying the presence of the virus in infected cells, characterization of neutralizing MAbs or antivirals. Thus, pH1N1-Nluc can be used to interrogate large libraries of biologics to identify those with neutralizing or antiviral activity for their potential use as therapeutics for viral infections. Our results are consistent with previous studies using reporter viruses for the identification of therapeutics against IAV (12, 14, 21, 48, 49).

Our *in vivo* experiments in B6 mice suggest that pH1N1-Nluc virus has similar virulence and similar levels of viral replication in NT and lungs, as pH1N1-WT virus. Importantly, Nluc activity in mice infected with pH1N1-Nluc was dose-dependent, with higher levels of Nluc activity in B6 mice infected with higher number of infectious viruses. Levels of Nluc expressions correlated with viral titers, providing an additional method to detect pH1N1 infection and to quantify the presence of the virus in infected mice. Importantly, expression of Nluc did not affect virus replication or pathogenicity in mice, contrary to viruses expressing fluorescent proteins (9, 12, 21, 22). Our initial studies in mice were further confirmed in ferrets, where we were able to detect viral infections in both infected and direct contact naïve animals at levels comparable to pH1N1-WT but with the advantage of being able to detect Nluc luminescence as a surrogate for viral infection. Our stability studies suggest that pH1N1-Nluc is more genetically and phenotypically stable than recombinant IAV expressing fluorescent proteins (9, 21, 25). The feasibility of implementing this approach to generate other IAV expressing Nluc from the NS segment was further demonstrated with the generation of a recombinant PR8-Nluc that retained similar replication efficiency and plaque phenotype as compared to PR8-WT in MDCK cells.

In summary, we describe the generation of a replication-competent recombinant pH1N1 virus expressing Nluc from a modified NS segment that is genetically and phenotypically stable and biologically comparable to pH1N1-WT virus both *in vitro* and *in vivo* for real-time tracking viral infection in living mice using IVIS. This Nluc expressing virus can serve as a robust platform for the identification of biologics with antiviral activity using HTS approaches. Moreover, we have also developed a similar Nluc-expressing PR8 virus, one of the most used and validated IAV in influenza research.

## MATERIALS AND METHODS

### Cells and Viruses

MDCK (Madin-Darby canine kidney) and human 293T cells were obtained from ATCC and grown in Dulbecco’s modified Eagle’s medium (DMEM: Gibco) supplemented with 5% fetal bovine serum (FBS) and 1% PSG (penicillin,100 units/ml: streptomycin 100UG/ml; L-Glutamine, 2mM) at 37°C with 5% CO_2_. The pH1N1-WT, pH1N1-Nluc, PR8-WT and PR8-Nluc were propagated in MDCK cells as previously described (50).

### Rescue of recombinant pH1N1-Nluc and PR8-Nluc viruses

Ambisence pDZ plasmids were used for the rescue of pH1N1-WT, pH1N1-Nluc, PR8-WT, and PR8-Nluc (51-53). Co-cultures (10^6^ cells/well) of 293T and MDCK cells (1:1) were co-transfected in suspension on 6-well plates with eight pH1N1 or PR8 ambisense pDZ plasmids (pDZ-PB2, -PB1, -PA, -HA, -NP, -NA, -M, -NS or -NS-Nluc). At 24 h after plasmid transfection, media was removed, and cells were incubated in DMEM with 0.3 % BSA and 1% PSG containing 0.5 µg/mL tosylsulfonyl phenylalanyl chloromethyl ketone (TPCK)-treated trypsin (Sigma). At 48 h post-transfection, media was collected and used to infect fresh monolayers of MDCK cells in 6-well plates (10^6^ cells/well). At 48 h post-infection, rescued viruses were confirmed by HA assay. Recovered viruses were plaque purified, and viral stocks were propagated in MDCK cells at 37°C in a 5% CO_2_ incubator for 3-4 days. Cell culture supernatants were collected, centrifuged ad 2,000 rpm for 5’ at room temperature, aliquoted, and storage at -80°C. For infections, virus stocks were diluted in PBS 0.3% BSA and 1% PSG. Following infection, cell monolayers were maintained in DMEM with 0.3% BSA and 1% PSG containing 0.5 µg/mL TPCK-treated trypsin. Viral titers (plaque forming units, PFU/mL) were calculated by standard plaque assay in MDCK cells as previously described (8, 43).

### Western blots

Cell extracts from mock or virus-infected (MOI=3) MDCK cells were collected and lysed in RIPA buffer. Proteins from lysates were separated by 12% SDS-PAGE, transferred to a nitrocellulose membrane, blocked in 5% fat-free dried milk in PBS containing 0.1% Tween 20 (PBS-T) and incubated overnight at 4°C with specific primary monoclonal (MAb) or polyclonal (PAb) antibodies against NS1, NP, or NLuc. Anti-glyceraldehyde 3-phosphate dehydrogenase (GAPDH) mAb (1:5000, Abcam, ab9484) was used as internal loading control. Bound primary antibodies were detected by HRP (horseradish peroxidase) conjugated secondary antibodies (1:2,000 dilution). Proteins were detected by chemiluminescence using a Molecular Imager Chemi Doc-XRS (Bio-Rad) (28).

### Virus growth kinetics

Multicycle growth kinetics were conducted in MDCK cells (12-well plate format, 5x10^5^ cells/well, triplicates). Briefly, subconfluent monolayers of MDCK cells were infected (MOI of 0.01) with the indicated viruses. After 1 h of viral adsorption, cell monolayers were overlayed with DMEM including 0.3% BSA, 1% PSG and 0.5 µg/mL TPCK-treated trypsin and incubated at 37°C. At 24, 48, 72 and 96 hpi, viral titers in cell culture supernatants were determined by immunofocus assay (fluorescent forming units, FFU/mL) using the NP MAb HB-65 as previously described (8). Nluc expression in the cell culture supernatants was quantified by using a Nano-Glo luciferase substrate (Promega).

### Plaque assays and immunostainings

Subconfluent monolayers of MDCK cells (6-well plate format, 10^6^ cells per well) were infected with the indicated viruses for 1 h at room temperature. After viral adsorption, virus inoculum was removed, and the cell monolayers were overlayed with infection media (DMEM 0.3% BSA, 1% PSG, 0.5 µg/mL TPCK-treated trypsin) containing agar. The plates were incubated at 37°C for 72 h. Then, plates were fixed overnight in 4% paraformaldehyde (PFA) and the overlay was removed. To visualize Nluc expression, plates were treated with Nano-Glo luciferase substrate and imaged under a Chemidoc (Bio-Rad). After detection of Nluc expression, plates were permeabilized with 0.5% Triton X-100 in PBS for 15 min and immunostained using the NP MAb HB-65. Immunostaining was developed using the vector kits (Vectastain ABC kit for mouse and DAB HRP substrate kit; Vector) following manufacturer specifications.

### Genetic stability of pH1N1-Nluc virus

MDCK cells (6-well plate format, 10^6^ cells per well) were infected (MOI of 0.01) with pH1N1-Nluc virus and incubated until 70% cytopathic effect (CPE) was observed. Cell culture supernatants were harvested and diluted (1:100), and similar infections of fresh MDCK cells (6-well plate format, 10^6^ cells per well) were followed for a total of 10 passages (P). Collected cell culture supernatants were also used to assess Nluc expression. Cell culture supernatants from P1, P5, and P10 were used to conduct plaque assays and evaluate Nluc expression and immunostained for NP using the MAb HB-65 (54).

### Nluc-based microneutralization assay for the identification of neutralizing antibodies

To test the neutralizing activity of MAb KPF1, confluent monolayers MDCK cells (96-plate format, 5x10^4^ cells/well, quadruplicates) were infected with 100 PFU of pH1N1-WT or pH1N1-Nluc for 1 h at 37°C. After viral adsorption, cells were washed and incubated with 100 μL of infection medium containing 2-fold serial dilutions (starting concentration of 0.25 μg/mL) of KPF1 or PBS, 1% Avicel (Sigma-Aldrich), and 1 μg/mL of TPCK trypsin. The plates were then incubated at 37°C for 24 h. Subsequently, the overlays were removed, and cells were fixed with 4% PFA for 1 h, washed with PBS and permeabilized with 100 μL of 0.5% Triton X-100 in PBS at room temperature for 15 min. Cells are then incubated with the mouse HB-65 primary antibody against NP for 1 h, followed by staining with an anti-mouse secondary antibody from the Vectastain ABC kit and a DAB peroxidase substrate kit (Vector Laboratories) according to the manufacturer’s recommendations. Viral plaques in each well were quantified using an ImmunoSpot plate reader. The viral titers were calculated as previously described (55-57). The mean ± SD of viral inhibitions are calculated from quadruplicate wells. Non-linear regression curves and IC_50_ values were determined using sigmoidal dose-response curves on GraphPad Prism.

For the Nluc activity-based microneutralization assay, MDCK cells were infected with 100 PFU of pH1N1-Nluc virus for 1 h. Cell monolayers were washed and incubated with 100 μL of infection media containing 2-fold serial dilutions (starting concentration of 0.25 μg/mL) of KPF1 or PBS, 1 μg/mL of TPCK trypsin, and incubated for 48 h. At 48 hpi, supernatants were collected, and Nluc activity was measured by adding Nluc substrate (1:50) at a 1:1 ratio. A Glowmax plate reader was used to read the Nluc activity in the cell culture supernatants. The mean ± SD of Nluc activity inhibitions were calculated from quadruplicate wells. Non-linear regression curves and IC_50_ values were determined using sigmoidal dose-response curves on GraphPad Prism.

### Nluc-based microneutralization assay for the identification of antivirals

To test the antiviral activity of Ribavirin, confluent monolayers of MDCK cells in 96 well plate (5×10^4^ cells/well, quadruplicates) were infected with 100 PFU of pH1N1-WT or pH1N1-Nluc for 1 h at 37°C. After viral adsorption, cells were washed and incubated with 100 μL of infection media containing 2-fold serial dilutions (starting concentration of 100 µM) of Ribavirin or PBS, 1% Avicel (Sigma-Aldrich), and 1 μg/ml of TPCK trypsin, and then incubated at 37°C for 24 h. Then, the overlay was removed, and cells were fixed with 4% PFA for 1 h. Subsequently, fixed cell monolayers were washed with PBS and permeabilized with 100µl of 0.5 % Triton X-100 in PBS at room temperature for 15 min. Cells were then incubated with the mouse HB-65 primary antibody against the viral NP for 1 h, followed by staining with an anti-mouse secondary antibody from the Vectastain ABC kit and a DAB peroxidase substrate kit (Vector Laboratories), according to the manufacturer’s recommendations. Viral plaques in each well were determined using an ImmunoSpot plate reader. The viral titers were calculated as previously described (55-57). The mean ± SD of viral inhibitions were calculated from quadruplicate wells. Non-linear regression curves and IC_50_ values were determined using sigmoidal dose-response curves on GraphPad Prism.

For the Nluc activity-based antiviral assay, MDCK cells were infected with 100 PFU of pH1N1-Nluc virus for 1 h. Cell monolayers were washed and incubated with 100 μL of infection media containing 2-fold serial dilutions (starting concentration of 100 μM) of Ribavirin or PBS, 1 μg/mL of TPCK trypsin, and incubated at 37°C in a 5% CO_2_ incubator. At 48 hpi, supernatants were collected, and Nluc activity was measured by adding Nluc substrate (1:50) at a 1:1 ratio. A Glowmax plate reader was used to read Nluc activity in the cell culture supernatants. The mean ± SD of Nluc activity were calculated from quadruplicate wells. Non-linear regression curves and IC_50_ values were determined using sigmoidal dose-response curves on GraphPad Prism.

### Mouse experiments

Six-week-old female C57BL/6J (B6) mice were purchased from the Jackson Laboratory (Maine, USA) and maintained in an animal facility at Texas Biomed under specific pathogen free conditions. For viral infection, cohorts of mice were anesthetized by gaseous sedation in an isoflurane chamber and intranasally inoculated with 10^2^, 10^3^, 10^4^, or 10^5^ PFU of pH1N1-WT or pH1N1-Nluc in a total volume of 50 µl (n=5 mice/group). Morbidity (changes in body weight) and mortality (% survival) of pH1N1-WT or pH1N1-Nluc in infected animals (n=5/group) were determined for 15 days. Mice with 25% weight loss from the initial body weight were determined as reaching the experimental end point and were humanely euthanized. Survival curves were plotted according to the Kaplan-Meier. Animals in the body weight group were also used to detect Nluc expression using an IVIS Spectrum multispectral imaging system. Briefly, mice were anesthetized on days 1, 2, 4, 6, 8 and 10 post-infection with isoflurane and injected with a 1:10 dilution in PBS of the Nano-Glo luciferase substrate retro-orbitally in a final volume of 100 µl and immediately imaged in the IVIS Spectrum multispectral imaging system. Data acquisition and analysis of Nluc expression in infected mice was conducted using the Aura program (AMI spectrum). Flux measurements were acquired from the region of interest. The scale used is indicated in each of the figures. Another group of mice were similarly infected and humanely euthanized at 2 and 4 dpi (n=3/group/time point). Before euthanasia, Nluc expression in the entire animals was determined as described above by anesthetizing the animals with isoflurane and retro-orbitally injection with 100 µL of Nano-Glo luciferase substrate (1:10). After imaging, lungs and nasal turbinate (NT) from naïve and infected mice were collected after euthanasia and homogenized in 1 mL of PBS using a Precellys tissue homogenizer (Bertin instruments) for 20 s at 7,000 rpm. Tissue homogenates were centrifuged at 12,000xg at 4°C for 5 min. Supernatants were collected and titrated by plaque assays and immunostaining as previously described (54, 58).

### Ferret experiments

To evaluate the feasibility of using pH1N1-Nluc to study virus pathogenicity and transmission in ferrets, outbred 4-month-old castrated male Fitch ferrets were obtained from Triple F Farms (Gillett, PA, USA). A contact transmission study was conducted with six ferrets per group: two ferrets were directly infected (10^6^ PFU/ferret) with either pH1N1-WT or pH1N1-Nluc and four naive ferrets were assigned as contacts (1:2 ratio). Briefly, on day 0, infected ferrets were anesthetized and inoculated intranasally with 10^6^ PFU of pH1N1-WT or pH1N1-Nluc and housed individually. At 24 hpi, two naive contact ferrets were introduced into each cage of the infected ferrets, resulting in three ferrets per cage (1 infected: 2 contact). Body weights were recorded at 0, 2, 4, 6, and 7 dpi. Nasal washes were collected from anesthetized ferrets on 2, 4, and 6 dpi to determine viral titers. On day 7, ferrets were humanely euthanized under anesthesia via exsanguination followed by intracardiac injection of Sleepaway euthanasia solution (Fort Dodge, Sodium Pentobarbital). Nasal turbinate (NT), olfactory lobes (OL), trachea, and lungs were collected for viral quantification by plaque assay and immunostaining as described above (54, 58).

### Genetic stability of pH1N1-Nluc *in vitro*

MDCK cells were (6-well plates, 10^6^ cells/well) were plated and infected (MOI of 0.01) with pH1N1-Nluc and incubated at 37°C in a 5% CO_2_ incubator until ∼70-80% cytopathic effect (CPE) was observed. Cell culture supernatants were then harvested and diluted (1:100) for subsequent serial passages in fresh MDCK cells (6-well plates, 10^6^ cells/well) for a total of 10 passages. At each passage, cell culture supernatants were used to assess Nluc expression and viral titration using plaque infectivity assay. Viral RNA from passages 1 (P1), P5 and P10 were extracted using TRIzol^TM^ reagent (Thermo fisher Scientific, US), for whole genome sequencing using next-generation sequencing platform MinION (Oxford Nanopore Technologies). Sample libraries were prepared using the Native Barcoding Kit 24 V14 (SQK-NBD114.24, Oxford Nanopore Technologies) and ran on R10.4.1. Flow Cells (FLO-MIN114, Oxford Nanopore Technologies) per manufacturer’s instructions. The read length stats with n50 v1.7.0 then the raw reads were trimmed using nanoq v0.10.0 (59). Bases with quality PHRED quality scores less than 7 were removed along with the first and last 25bp of each read. Reads less than 500 bp after trimming were removed from downstream analyses. Filtered reads were mapped to the HPhTX NSs-Nluc reference sequence using minimap2 v2.28-r1209 (60) and the ‘-x map-ont’ option for mapping long error prone nanopore reads. Mapping rates were calculated with SAMtools flagstat v1.21 (61) and reads with mapping quality (MAPQ) scores less than 40 were removed as well as reads mapping to less than 1,000 bp of the reference. We estimated coverage across each genome with MosDepth v0.3.10 (62). We reevaluated indel quality scores and called variants using LoFreq v2.1.5 (63). We limited the depth of coverage (--max-depth) for variant calling at 5,000x and removed the default filters applied by LoFreq. We used custom filters to remove any variants that were present in less than 25% of reads, with read depths less than 100x, or were in homopolymer regions of 3bp or larger.

### Ethical Approvals

The Institutional Animal Care and Use Committee (IACUC) of Texas Biomed approved all mouse experiments conducted in this study (IACUC# 1785MU). Ferret experiments were performed in accordance with the animal protocol approved by the IACUC of the Icahn School of Medicine at Mount Sinai (ISMMS) (IACUC# IACUC-2013-1408). All mice and ferrets were housed in a temperature- and humidity-controlled Animal Biosafety Level 2 (ABSL-2) facility. All procedures throughout the study were designed to minimize animal suffering. The Institutional Biosafety Committees (IBC) of Texas Biomed and the ISMMS reviewed and approved of the generation and use of genetically engineered influenza viruses.

### Quantifications and statistical analyses

All graphs, calculations, and statistical analyses were performed using GraphPad Prism software version 9.5.1 (GraphPad Software, LLC, USA). Growth kinetics were analyzed using a two-way repeated measure ANOVA with Geisser-Greenhouse correction. Post-hoc comparisons were performed using the Šídák method. Differences in FFluc expression were evaluated using Welch’s one-way ANOVA, followed by multiple comparisons using Dunnett T3 method. Viral replication and titer data were analyzed using mixed-effects ANOVA. Dunnett’s multiple comparisons test was used to compare groups within each time-point. Post-hoc testing was conducted with Dunnett’s multiple comparisons test. The significance of differences are indicated as follows: ns=non-significant, * = p < 0.05, ** = p < 0.01, *** = p < 0.001, **** = p < 0.0001.

## DECLARATIONS

### Data availability

All NGS raw data are available under the BioProject PRJNA1288407 (https://www.ncbi.nlm.nih.gov/bioproject/PRJNA1288407).

## AUTHOR CONTRIBUTIONS

Conceptualization: R.S.B., A.M., A.N., and L.M-S.; Methodology: R.S.B., A.M., K.C., R.L.P., R.N.P., A.C., R.A.A., A.N., and L.M-S.; Data collection and interpretation: R.S.B., A.M., A.A-G., T.J.C.A., U.G.K., R.A.A., A.G-S, J.J.K., A.N., and L.M-S.; Funding acquisition and resources: A.G-S., U.G.K., R.A.A., A.G-S., J.J.K. and L.M-S.; Writing— original draft preparation: R.S.B., A.M. and L.M-S.; Writing—review and editing: all authors have read and agreed to the published version of the manuscript.

