## Supplemental Figures for "Bioluminescent reporter influenza A viruses to track viral infections"

<table>
<tr><th>Gene</th><th>Coverage</th><th>SNP(S)</th></tr>
<tr><td>PB2</td><td>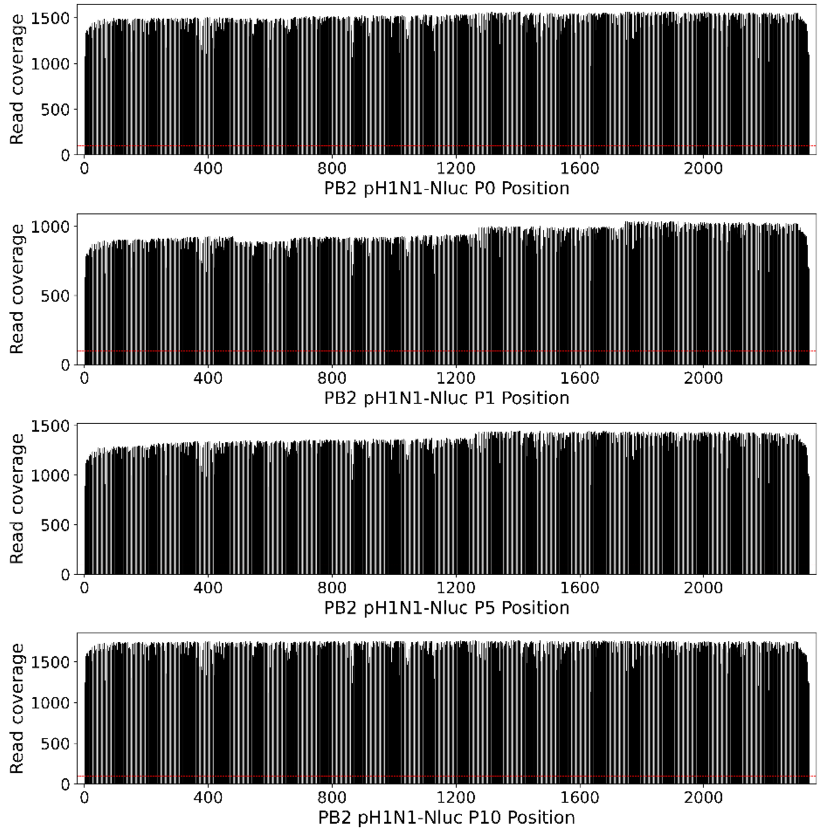
</td><td>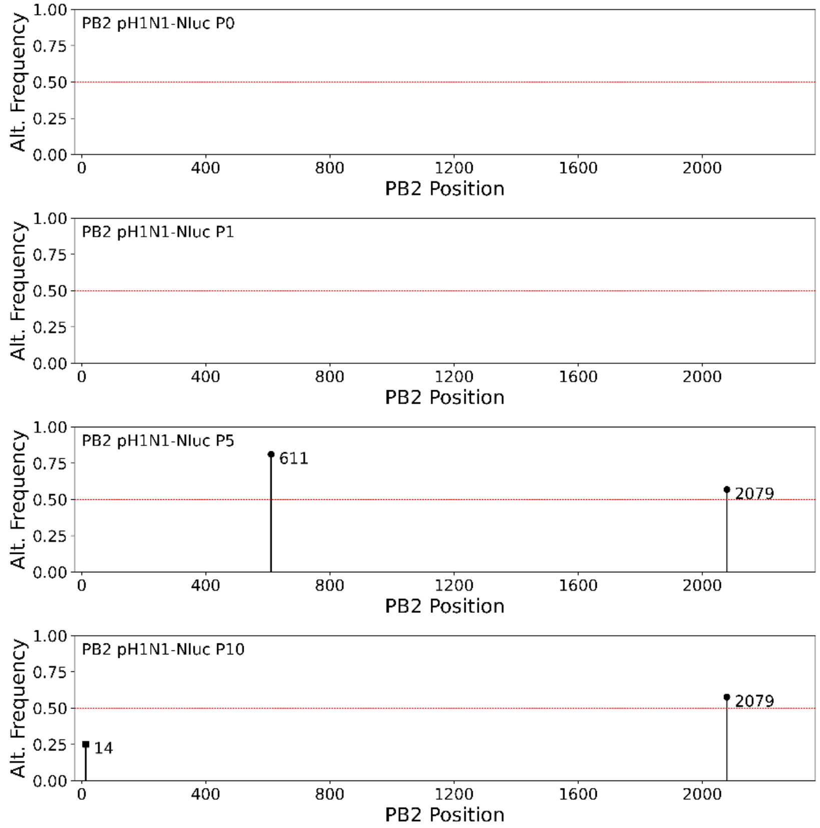
</td></tr>
</table>

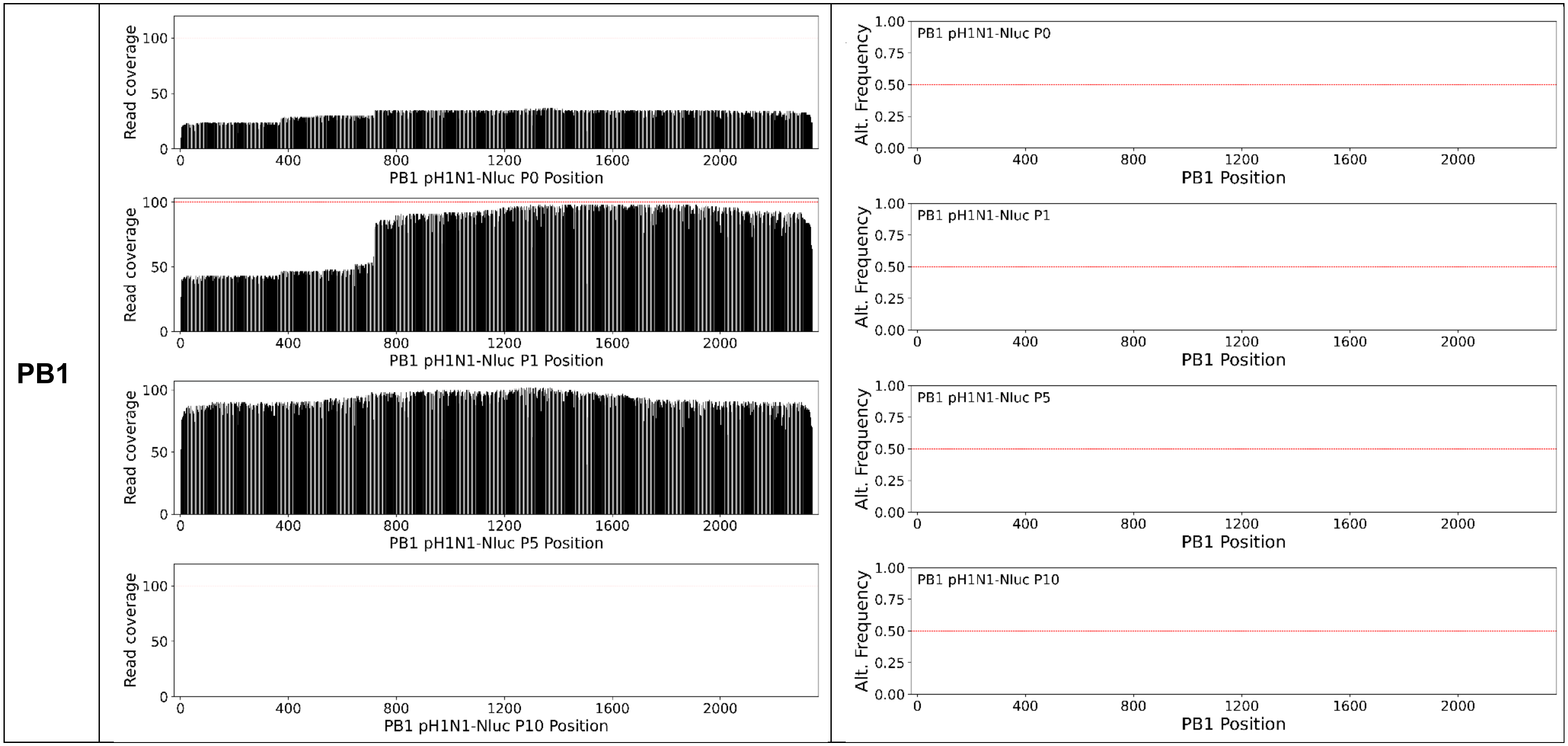
PB1
Read coverage
PB1 pH1N1-Nluc P0 Position
PB1 pH1N1-Nluc P1 Position
PB1 pH1N1-Nluc P5 Position
PB1 pH1N1-Nluc P10 Position
Alt. Frequency
PB1 pH1N1-Nluc P0
PB1 pH1N1-Nluc P1
PB1 pH1N1-Nluc P5
PB1 pH1N1-Nluc P10
PB1 Position

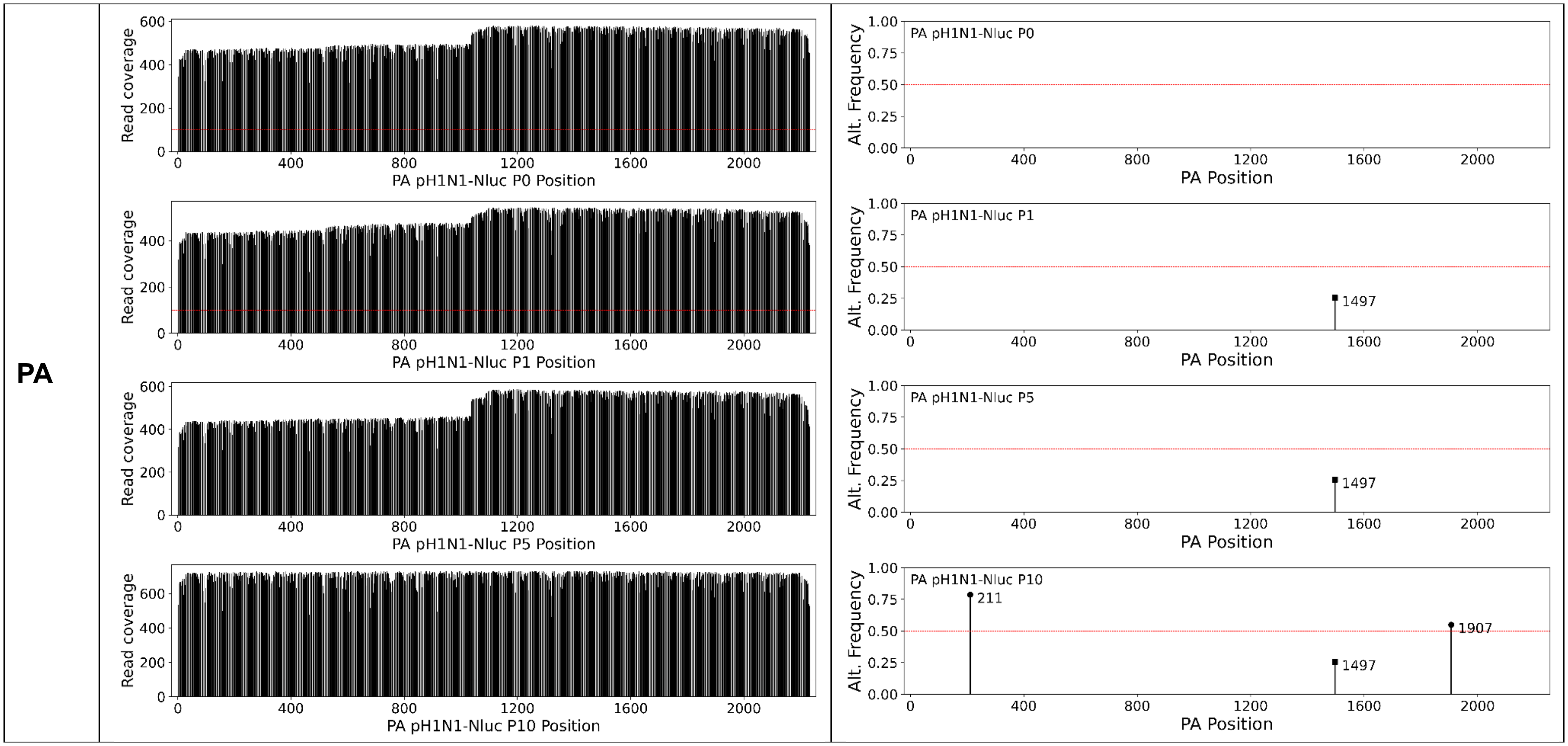
PA
Read coverage
PA pH1N1-Nluc P0 Position
PA pH1N1-Nluc P1 Position
PA pH1N1-Nluc P5 Position
PA pH1N1-Nluc P10 Position
Alt. Frequency
PA Position
PA pH1N1-Nluc P0
PA pH1N1-Nluc P1
1497
PA pH1N1-Nluc P5
1497
PA pH1N1-Nluc P10
211
1907
1497

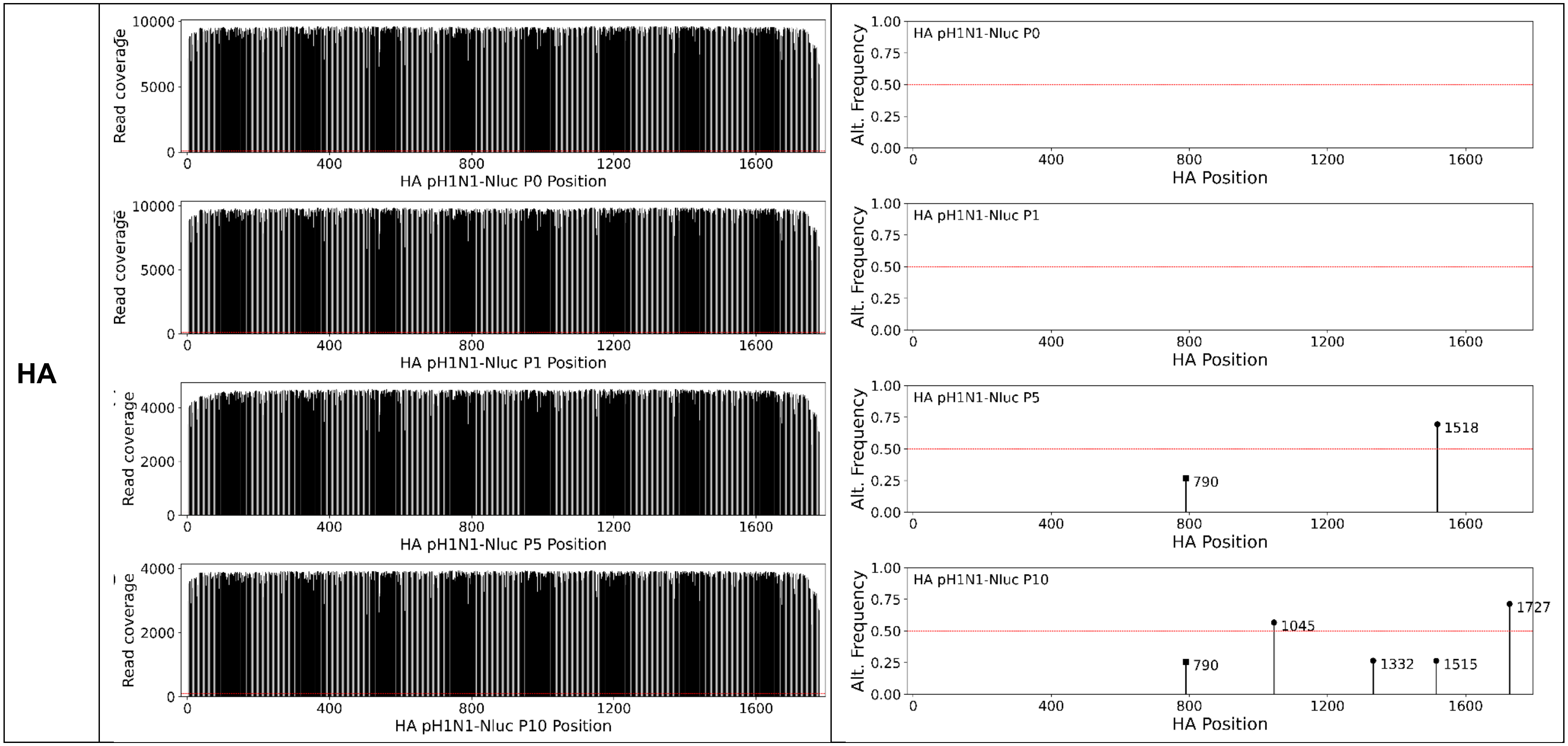

HA
Read coverage
HA pH1N1-Nluc P0 Position
HA pH1N1-Nluc P1 Position
HA pH1N1-Nluc P5 Position
HA pH1N1-Nluc P10 Position
Alt. Frequency
HA Position
HA pH1N1-Nluc P0
HA pH1N1-Nluc P1
HA pH1N1-Nluc P5
790
1518
HA pH1N1-Nluc P10
790
1045
1332
1515
1727

NP

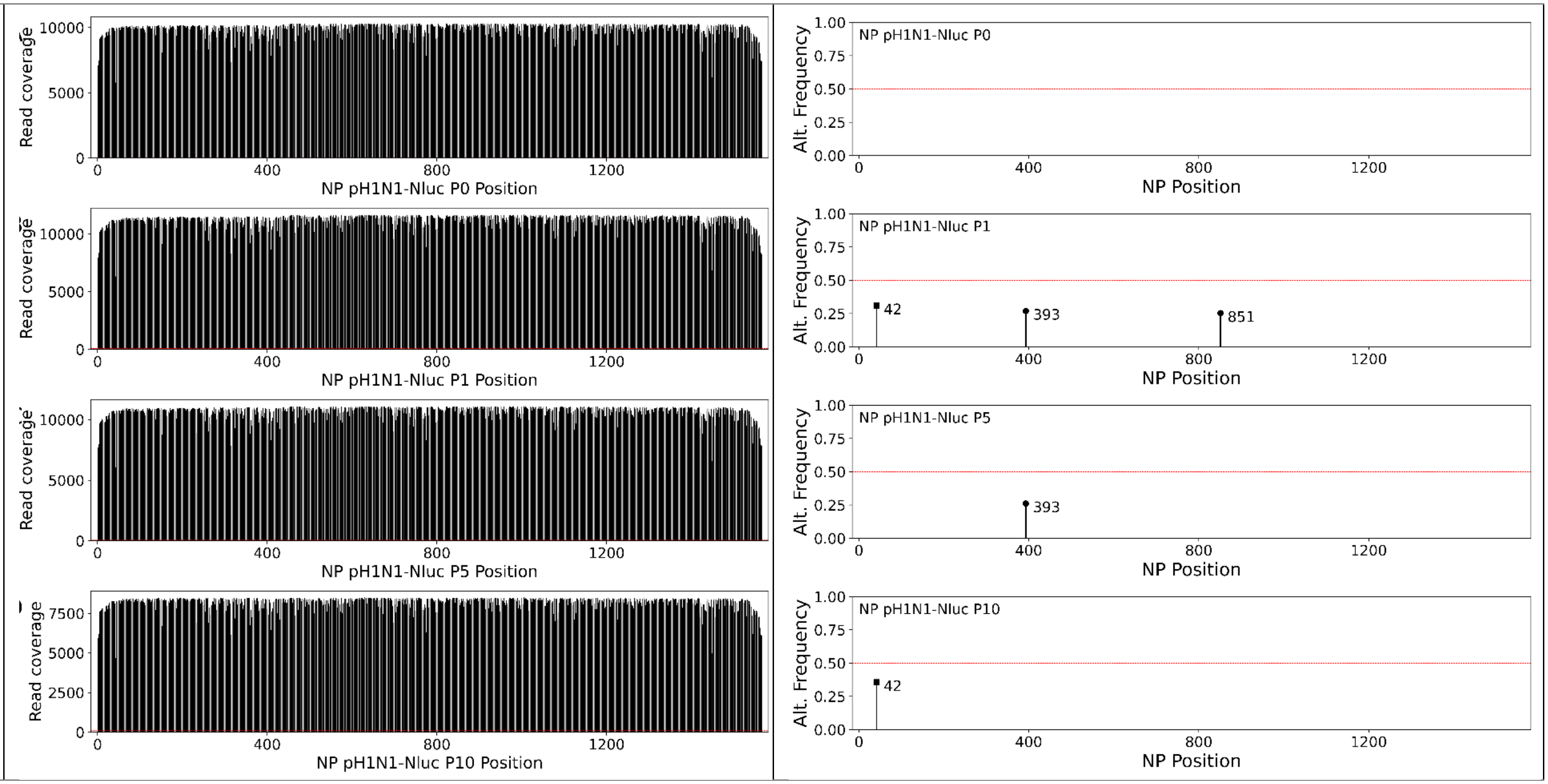

NP pH1N1-Nluc P0 Position
NP pH1N1-Nluc P1 Position
NP pH1N1-Nluc P5 Position
NP pH1N1-Nluc P10 Position
Read coverage
NP pH1N1-Nluc P0
NP pH1N1-Nluc P1
NP pH1N1-Nluc P5
NP pH1N1-Nluc P10
Alt. Frequency
NP Position
42
393
851
393
42

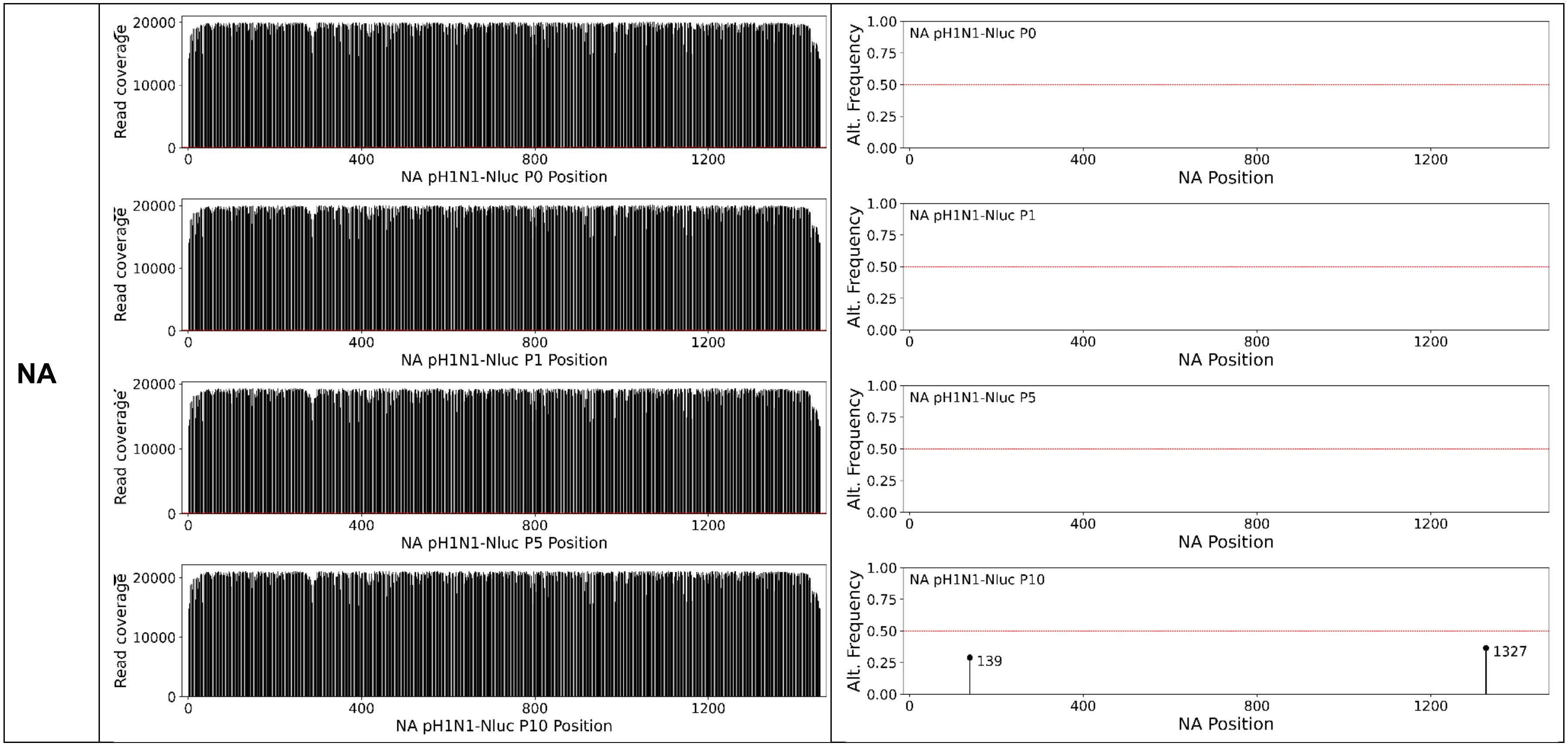

NA
Read coverage
20000
10000
0
0
400
800
1200
NA pH1N1-Nluc P0 Position
NA pH1N1-Nluc P1 Position
NA pH1N1-Nluc P5 Position
NA pH1N1-Nluc P10 Position
Alt. Frequency
1.00
0.75
0.50
0.25
0.00
NA pH1N1-Nluc P0
NA pH1N1-Nluc P1
NA pH1N1-Nluc P5
NA pH1N1-Nluc P10
139
1327
NA Position

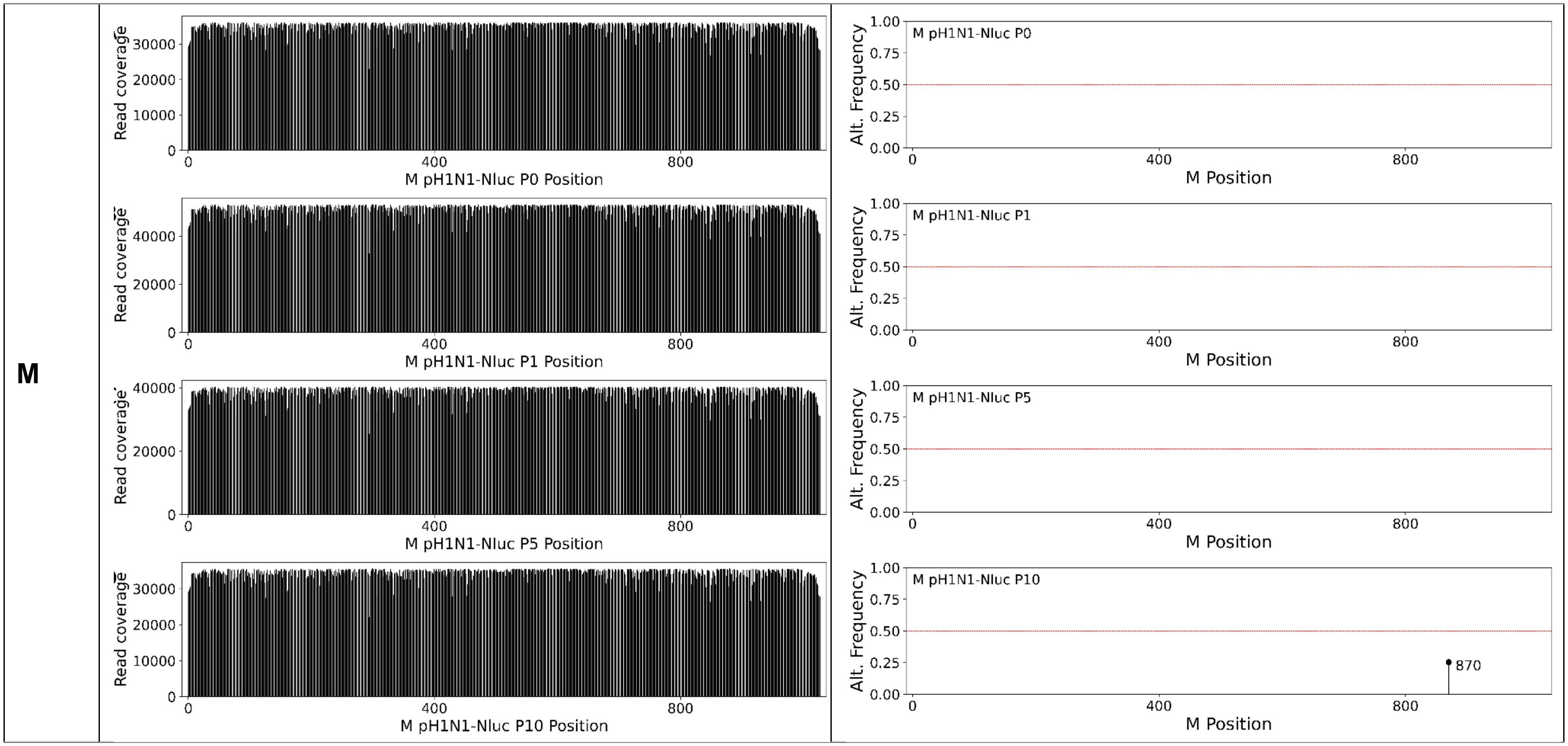


**Supplementary Figure S1. Genetic stability of pH1N1-Nluc viral segments PB2, PB1, PA, HA, NP, NA, and M following *in vitro* passaging. SNPs: Single nucleotide polymorphisms.**

**Supplementary table 1.** ***Data description*** **-** Sample summary and genome coverage

| Virus_passage | PE reads (1e4) | Alignment Rate | Mean Coverage |
|---|---|---|---|
| pH1N1-Nluc_P0 | 9 | 99.98% | 8029.3 |
| pH1N1-Nluc_P1 | 131.4 | 99.89% | 9251.6 |
| pH1N1-Nluc_P5 | 145.5 | 99.50% | 7761.8 |
| pH1N1-Nluc_P10 | 116.3 | 99.81% | 8670 |
